# Dispersal provides trophic-level dependent insurance against a heatwave in freshwater ecosystems

**DOI:** 10.1101/2022.09.20.508571

**Authors:** Csaba F. Vad, Anett Hanny-Endrédi, Pavel Kratina, András Abonyi, Ekaterina Mironova, David S. Murray, Larysa Samchyshyna, Ioannis Tsakalakis, Evangelia Smeti, Sofie Spatharis, Hanrong Tan, Christian Preiler, Adam Petrusek, Mia M. Bengtsson, Robert Ptacnik

## Abstract

Climate change-related heatwaves are major recent threats to biodiversity and ecosystem functioning. However, our current understanding of the mechanisms governing community resilience (resistance and recovery) to extreme temperature events is still rudimentary. The spatial insurance hypothesis postulates that diverse regional species pools can buffer ecosystem functioning against local disturbances through immigration of better adapted taxa. However, experimental evidence for such predictions from multi-trophic communities and pulse-type disturbances, like heatwaves, are largely missing. We performed an experimental mesocosm study with alpine lake plankton to test whether a dispersal event from natural lakes prior to a simulated heatwave could increase resistance and recovery of local communities. As the buffering effect of dispersal may differ among trophic groups, we independently manipulated dispersal of organisms from lower (microorganisms) and higher (zooplankton) trophic levels. The experimental heatwave suppressed total community biomass by having a strong negative effect on zooplankton biomass, probably due to a heat-induced increase in metabolic costs that in turn caused mortality. Heating thus resulted in weaker top-down control and a subsequent shift to bottom-heavy food webs. While zooplankton dispersal did not alleviate the negative heatwave effects on zooplankton biomass, dispersal of microorganism enhanced biomass recovery at the level of phytoplankton, thereby providing evidence for spatial insurance. The different response of trophic groups may be related to the timing of dispersal, which happened under strongly monopolized resource conditions by zooplankton, creating limited opportunity for competitors to establish. At the same time, the heatwave released phytoplankton from grazing pressure and increased nutrient recycling, which may have facilitated the establishment of new phytoplankton taxa. Our findings clearly show that even a short heatwave can strongly alter energy flow in aquatic ecosystems. Although dispersal can enhance community resilience, the strength of its buffering effects depends on the trophic level.

## 1. Introduction

Global climate change is characterised not only by rising means in annual surface temperatures but also by increasing frequency, magnitude and duration of heatwaves (IPCC, 2021). There is evidence for these increasing trends in heatwaves across the terrestrial (Fischer & Schär, 2010; Perkins-Kirkpatrick & Lewis, 2020), marine (Frölicher et al., 2018; Oliver et al., 2018), and freshwater realms (Woolway et al., 2021). Although the critical role of extreme whether events driving ecosystem changes has long been recognized (Jentsch et al., 2007), much of the previous climate change research has focused on the effects that rising mean temperatures. For example, a large majority of experimental studies aiming to unravel ecosystem responses to temperature increases applied static warming treatments without incorporating extreme events (Thompson et al., 2013; Woodward et al., 2016). Consequently, our understanding of how ecological communities and ecosystems respond to extreme weather events, such as heatwaves, is still limited.

Heatwaves often impose short but intense disturbances. By quickly pushing organisms beyond their thermal tolerance limits and compromising their physiological or genetic adaptations, heatwaves may alter community composition and ecosystem functioning more strongly than a gradual rise in mean temperatures (Bennett et al., 2021; Gutschick & BassiriRad, 2003; Stillman, 2019; Vasseur et al., 2014). It has been suggested that frequent heatwaves can reshuffle global biodiversity patterns by causing local extinctions coupled with species range shifts (Smale & Wernberg, 2013; Wernberg et al., 2013), modulating population dynamics (Davison et al., 2010; Jiguet et al., 2006), and altering species interactions (Sentis et al., 2013; Zhang et al., 2018). All these changes can in turn impair ecosystem functioning (Eggers et al., 2012; Thompson et al., 2015) and the provisioning of ecosystem services (Smale et al., 2019). Aquatic ecosystems may be particularly susceptible to heatwaves as aquatic ectotherms tend to exhibit narrower thermal safety margins than terrestrial ones (Pinsky et al., 2019; Sunday et al., 2012). Higher sensitivities to warming imply more frequent extinctions and faster species turnover in aquatic ecosystems, with implications for ecosystem functioning (Comte & Olden, 2017; Pinsky et al., 2019). However, in contrast to a press disturbance of steadily rising mean temperatures, a short-term pulse disturbance caused by a heatwave is likely to be followed by a certain degree of community and ecosystem recovery (Bender et al., 1984; Harris et al., 2018). Management strategies will therefore critically depend on our understanding of the mechanisms that govern the resilience of ecosystems against heatwaves, in particular of its key components, resistance to and recovery from a disturbance (Hodgson et al., 2015; Ingrisch & Bahn, 2018).

According to the spatial insurance hypothesis, the resilience of local communities to disturbance likely depends on the connectivity to and diversity of the surrounding regional species pool (Loreau et al., 2003; Thompson et al., 2017). This implies that habitats that are geographically isolated, either naturally or through anthropogenic impacts (e.g., habitat fragmentation), are likely to be more susceptible to environmental change, including more frequent heatwaves. Immigration of species more tolerant to certain disturbances may allow better tracking of the changing environment, allowing for more stable ecosystem functioning if the colonising and resident species are redundant in maintaining specific ecosystem processes (Loreau et al., 2003). However, experimental evidence in support of the spatial insurance hypothesis is still contradictory and no consensus has been reached. This is partly due to the fact that the insurance effect depends on the type of stressor and the measure of ecosystem functioning (Symons & Arnott, 2013; Thompson & Shurin, 2012). The spatial insurance can also differ among trophic groups due to their different responses to environmental stressors and abilities to disperse (Limberger et al., 2019). Yet, most previous experiments have focused only on simplified ecosystems composed of a single trophic group (de Boer et al., 2014; Eggers et al., 2012; Guelzow et al., 2017), or manipulated dispersal of only a single trophic group (Symons & Arnott, 2013; Thompson & Shurin, 2012). It has also been recently debated whether dispersal can provide spatial insurance against heatwaves. Laboratory experimental manipulations of a single trophic level suggested either positive (de Boer et al., 2014) or neutral (Eggers et al., 2012). As dispersal of organisms from different trophic levels differs in their effect on metacommunity structure and ecosystem function (Haegeman & Loreau, 2014), the direct experimental manipulation of multiple trophic levels in a metacommunity context can provide a more realistic understanding of ecosystem resistance to and recovery from extreme heatwaves on a regional scale.

There has been mounting evidence that the impacts of warming critically depend on the trophic level (and associated traits such as body size) of organisms, driven by their different physiological constraints and changes in the strength of trophic interactions (Kratina et al., 2022; Petchey et al., 1999; Shurin et al., 2012). For instance, increased metabolic demands of ectothermic consumers can result in higher feeding rates, resulting in stronger top-down control (Brown et al., 2004; Romero et al., 2018; P. Zhang et al., 2020). At the same time, large consumers are more prone to starvation under warmer conditions, which increases the risk of local extinction (Fussmann et al., 2014; Rall et al., 2010). It is further accentuated by their generally smaller population sizes and slower growth rates (Petchey et al., 1999; Purvis et al., 2000). Moreover, the successful establishment of consumer populations in a new habitat strongly depends on the availability of resources (Thompson & Gonzalez, 2017). Therefore, consumers may be more dispersal limited than their resources, resulting in stronger and longer-lasting responses to and slower recovery following disturbance. Lastly, it is essential to partition dispersal and associated diversity changes at different trophic levels, as responses of ecosystem functioning (e.g., primary production) to diversity changes directly depend on which trophic group is being affected (Duffy et al., 2007; Thébault & Loreau, 2003).

Here we tested how multi-trophic aquatic communities respond to heatwaves, and whether the spatial insurance effect of dispersal modulates community responses. We performed a mesocosm experiment where we first established plankton communities from a geographically isolated alpine lake. We then tested whether an initial dispersal event from a diverse regional species pool contributed to the resistance and recovery of the experimental communities during and after the heatwave manipulation. To be able to partition dispersal effects between trophic levels, dispersal was manipulated separately for microorganisms and zooplankton. We first hypothesised that disturbance caused by the experimental heatwave would result in reduced total community biomass. Second, given the different metabolic constraints and sensitivity to resource availability, we predicted that organisms at higher trophic levels (i.e., zooplankton) would be more negatively affected than those at lower positions, resulting in weaker top-down control. Thirdly, we hypothesised that increased connectivity to a regional species pool would enhance the community resistance to and recovery following the heatwave and this spatial insurance effect would have a higher importance for organisms at higher trophic levels.

## 2. Materials and methods

### 2.1 Experimental setup

We performed an outdoor mesocosm experiment between June and August 2018 at the Biological Station of WasserCluster Lunz, Austria. We investigated the independent and interactive effects of heatwave and dispersal on community composition and ecosystem functioning in a full-factorial design. Presence (H+) or absence (H-) of the heatwave was crossed with the manipulation of dispersal, represented by a dispersal event from the regional species pool of natural lakes and applied separately for microorganisms (M-, M+) and zooplankton (Z-, Z+). The experimental setup thus comprised 8 treatments and 5 replicates per treatment for a total of 40 experimental units.

The experimental system consisted of 40 land-based mesocosms (height: 81.0 cm, inner diameter: 77.0 cm) made of food-safe PE containers (ARICON Kunststoffwerk GmbH, Solingen, Germany). We placed them on an unshaded meadow approximately 500 m from Lake Lunz, Eastern Alps (N 47°51’15’’ E 15°03’07’’, 608 m a.s.l). Each mesocosm was insulated with mineral wool and covered by white opaque plastic foil on the outer side to reduce the thermal impact of air temperature and irradiation. As a result, the average diurnal fluctuations in mesocosm water temperatures were in the range commonly seen in the surface water of Lake Lunz. We covered the mesocosms with 250 μm-mesh net lids to minimise introduction of particles while allowing for air exchange.

At the start of the experiment, we filled the mesocosms with 300 L of water (resulting in a water depth of 66.6 cm) from Lake Lunz. Water was collected from a 2-m depth (i.e., from the lake epilimnion) by a centrifugal pump, transported by a water truck to the experimental site, and randomly pumped into the mesocosms after passing through a coarse sieve (500 μm) to exclude fish larvae. As mesozooplankton (especially cladocerans) were impaired by pumping, we also introduced natural lake zooplankton from net hauls, to set a starting density of ca. 3 *Daphnia* individuals L^-1^ in the mesocosms. This density corresponded to the mean summer density of *Daphnia* in natural lakes in the area (Horváth et al., 2017). In the oligotrophic (5–8 μg total phosphorus L^−1^) Lake Lunz, phosphorus is the limiting nutrient for primary production. After filling the mesocosms with lake water and organisms, total phosphorus (TP) concentrations were raised to 15 μg L^-1^ by addition of K_2_HPO_4_, to set slightly mesotrophic conditions, as we expected a reduction of nutrients through sedimentation over the course of the experiment. No experimental treatments were applied for 8 days, allowing for local species sorting and community establishment. To minimise the growth of periphyton and its impact on the planktonic system, we turned the removable inner wall and bottom plates of the mesocosms every other week.

To simulate dispersal from the regional species pool, we introduced a pooled inoculum consisting of either microorganisms (M+ treatment), zooplankton (Z+ treatment), or both (M+Z+ treatment), originating from 15 regional lakes (Table S1). Among these source lakes, we also included peri-alpine lowland habitats that more likely contain heat-tolerant plankton taxa compared to our focal site. To apply the treatments, we first collected samples of microorganisms and zooplankton from each source lake. For microorganisms, we collected a 20-L vertically integrated epilimnetic water sample with a Van Dorn bottle. In the next step, samples were pooled and screened through a plankton net (mesh size: 30 μm) to remove larger organisms, especially metazoans. Thus, this pooled sample contained all microorganisms <30 μm (i.e., viruses, bacteria and protists). We then introduced a 3-L subsample (representing 1% of total mesocosm volume) to inoculate the M+ mesocosms (N = 20). We collected zooplankton by vertical net hauls (mesh size: 100 μm, opening diameter: 40 cm) from the epilimnion of the 15 source lakes. These samples were first subsampled to the same effective volume per lake (20 L) as in the case of microorganisms. Before applying the Z+ dispersal treatment, the pooled community was washed gently by retaining zooplankton in a 100-μm-mesh plankton net that was kept submerged in a large bucket just below the rim while gently pouring water in. This way we ensured the animals did not fall dry, while most bacteria and protists were washed out. We then added a zooplankton dispersal inoculum corresponding to 3 L of lake water to each Z+ mesocosm. By this we introduced on average 160 individuals of cladocerans (corresponding to a density of ∼0.5 ind L^-1^ in the mesocosms) and 412 (∼1.4 ind L^-1^) individuals of copepods into each Z+ mesocosms, including 4 cladoceran and 4 copepod species that were not present in the local community (Table S2). Altogether, these steps allowed us to introduce an inoculum of standardised volume independently for the microorganism (M+) and zooplankton (Z+) dispersal treatments. The dispersal event (M+ and Z+ treatments) was simulated on day 8 of the experiment, prior to the experimental heatwave which started 48 h later (i.e., on day 10, Figure 1).

**Figure 1.**
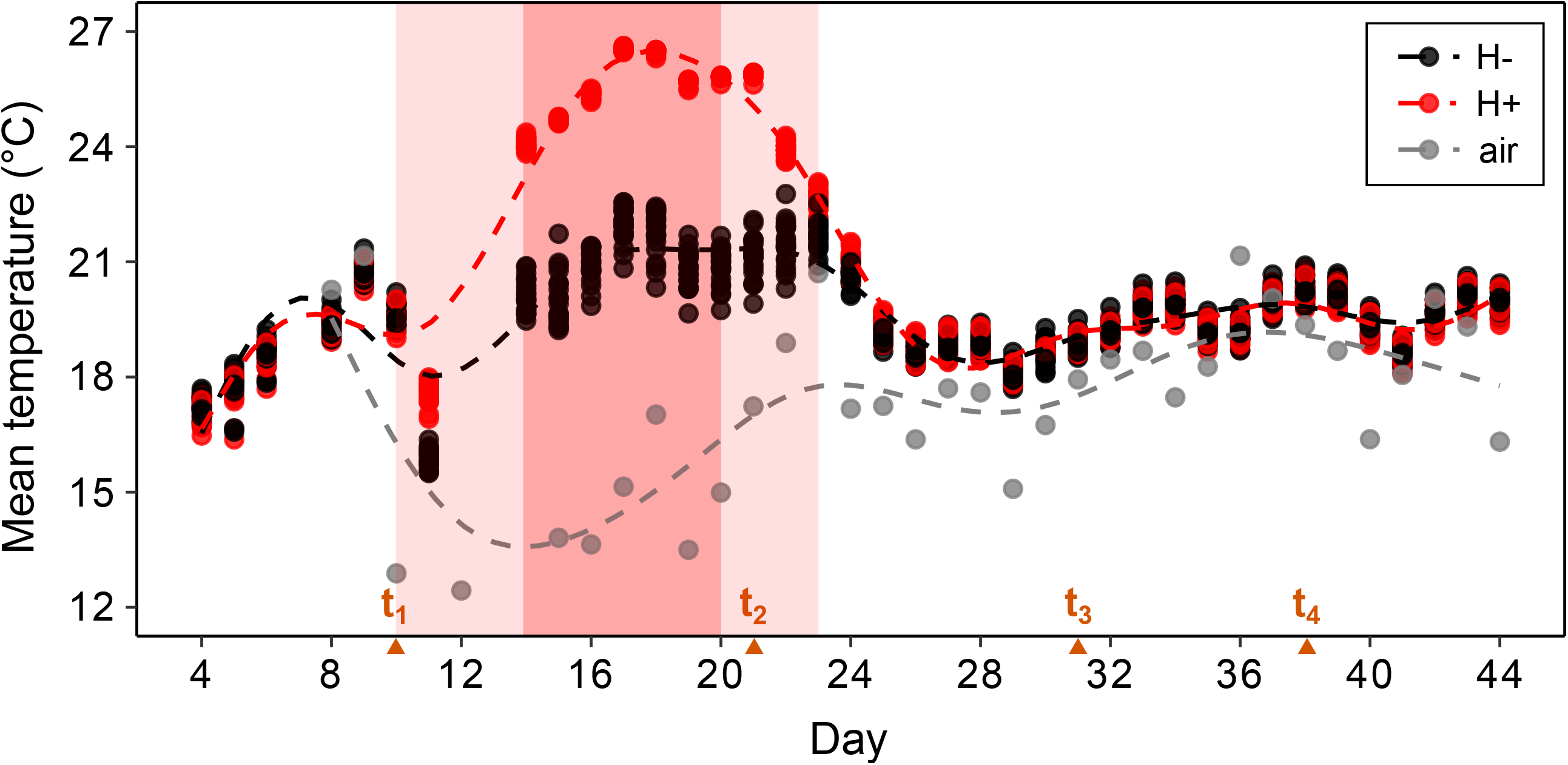
Temporal dynamics of daily mean water temperature and ambient air temperature over the experimental duration. Heatwave treatment (H+) is indicated with red, while control (H-) mesocosms are indicated with black (N = 20 per treatment and per day). Red-coloured shading denotes the time interval of the simulated heatwave in H+ treatments (light shading: heating and cooling phases, dark shading: culmination phase with a +5°C offset in H+ vs H-). Triangles show the timing of the four focal sampling campaigns (i.e., t_1_, t_2_, t_3_, and t_4_). Dashed lines represent fitted GAM models illustrating temperature dynamics.

The experimental heatwave (H+ treatment) was simulated by gradually increasing temperatures to reach a 5°C offset (required 4 days of initial heating) compared to the control (H-) conditions, which was subsequently maintained for 7 days (culmination phase). While we planned to use the ambient temperatures as baseline, due to unusually cold weather conditions, H-tanks were also heated from the third day of the heatwave to maintain a baseline temperature of 21.0°C until the end of the heating in H+ tanks. After turning off the heating, water temperature in H+ tanks returned to ambient levels within three days. The heatwave, including its initial heating, culmination, and cooling phases, lasted 14 days (Figure 1), and was followed by a 21-days recovery phase. The entire experiment (including the establishment, heatwave and recovery phases) therefore lasted for 44 days (Figure 1). During the heatwave, temperature was regulated by submersible 200-W aquarium heaters (thermocontrol 200, Eheim GmbH, Deizisau, Germany) connected to a computer-controlled heating system. To set temperature levels in the H+ treatment, the mean temperature of the control (H-) mesocosms (N = 20) was used as a baseline. To prevent the development of vertical gradients in the tanks we applied an airlift system (Striebel et al., 2013). Compressed air released from a tube produced a very gentle upward current in a PVC pipe hanging in the centre of each mesocosm, and this kept the entire water column constantly mixed during the experiment.

### 2.2. Sampling and sample processing

In vivo chlorophyll *a* (Chl-*a*) autofluorescence (hereinafter referred as Chl-*a* fluorescence) was measured daily by a handheld fluorometer (AquaPen-C AP-C 100, PSI, Drásov, Czech Republic) after a 30-minute dark-adaptation period, and served as a proxy for phytoplankton biomass. Samples were taken from the central surface water of the mesocosms. Over the experimental duration, samples were collected twice per week for TP, Chl-*a* (based on pigment extraction), particulate organic carbon (POC) as well as phyto- and zooplankton. Microscopic analysis of plankton community samples was carried out on four focal sampling dates: (i) two days after the introduction of the regional inoculum, but before starting the heatwave manipulation (day 10, t_1_), (ii) at the end of the culmination phase of the experimental heatwave when the heaters were turned off (day 21, t_2_), as well as (iii) 10 days (day 31, t_3_) and (iv) 17 days (day 38, t_4_) later, in the recovery phase (Figure 1). These dates were specifically chosen to test for resistance to the heatwave (i.e., the comparison between t_1_ and t_2_), and recovery during the post-heatwave recovery phase (i.e., the comparison between t_3_ and t_4_; see more details in Data analysis). Samples were collected through a tap at the side of each mesocosm (inner diameter: 10.0 cm, height from ground: 50.0 cm), to reduce the risk of unintentional dispersal (e.g., by a sampling device) among the experimental units. Prior to the sampling, we increased airflow and gently mixed the water column of each mesocosm with a clean plastic tube, ensuring a homogenous distribution of plankton. We then collected zooplankton samples by releasing 20 L from the tap into a clean container, thereby ensuring fast flow that prevents the zooplankton from escaping the suction. Subsequently, we filtered the volume through a 30-μm mesh plankton net and preserved the retained organisms in absolute ethanol. To obtain samples of phytoplankton, Chl-*a*, POC, and water for nutrient analysis, another 3-L water sample was collected and filtered through a 100-μm mesh to remove large zooplankton. For phytoplankton samples, 200 mL of water was preserved with Lugol’s iodine solution. 500 mL of water were filtered through glass microfiber filters for analysis of Chl-*a* and POC (Whatman GF/F, pore size: 0.7 μm), and filters were kept frozen (−20°C) until analysis. We then replaced the sampled water volume in each mesocosm with sterile-filtered (polyethersulfone membrane, pore size: 0.2 μm, MTS & APIC Filter, Bad Liebenzell, Germany), chlorine-free tap water, and added 15 μg L^-1^ K_2_HPO_4_ corresponding to the exchanged volume of water. As a result, TP concentrations generally varied between 10 and 15 μg L^-1^ over the experimental period and we did not find any systematic deviations across treatments (Figure S1).

Chl-*a* concentration was determined by fluorometry after acetone extraction (Arar & Collins, 1997), without correcting for phaeophytin. POC content was measured by an elemental analyser (vario MICRO cube™, Elementar Analysensysteme GmbH, Langenselbold, Germany). Concentration of TP was measured by the ascorbic acid colorimetric method (Hansen & Koroleff, 1999) after persulfate digestion (Clesceri et al., 1999).

### 2.3. Microscopic analyses

We estimated densities in the phytoplankton samples by the Utermöhl (1958) method with an inverted microscope (DMI3000 B, Leica Microsystems, Wetzlar, Germany). We counted and identified (to species level when possible) at least 400 sedimentation units (e.g., filaments, colonies, or single cells) in each sample (Lund et al., 1958). To obtain taxon-specific biovolume and wet weight, we applied conversion factors for corresponding geometrical shapes (Hillebrand et al., 1999), based on measurements of axial dimensions of at least 30 individuals for dominant taxa. We then converted wet weight to carbon mass by a conversion factor of 14% (Vadstein et al., 1988).

To obtain zooplankton density data, we counted all crustacean individuals present in the 20-L samples. For rotifers, we counted all individuals in subsamples representing 10% of the sampled volume. Individuals were identified to species level when possible. Specimens belonging to the *D. longispina* species complex were pooled, due to difficulty of reliable phenotypic differentiation between parental taxa and interspecific hybrids during routine identification (Dlouhá et al., 2010). We determined crustacean zooplankton body size by measuring the length of the first 20 individuals of each of the dominant species using a stereo microscope (Stemi 2000-C, Carl Zeiss AG, Jena, Germany), while we used published average length data for rotifers (Koste, 1978). Body length of cladocerans was measured from the top of the head to the base of the caudal spine; the length of copepods was measured from the tip of the cephalothorax to the base of the furca (Bottrell et al., 1976). For rare crustacean species, we used mean body size measurements obtained from replicates of the same treatment combination with higher densities. When a species occurred at low densities in all replicates of a treatment combination, mean body size was obtained from all the individuals available. For copepod nauplii, we used the published mean body length of *Cyclops abyssorum* nauplii, the dominant species in our experiment (Ludovisi et al., 2008). We subsequently converted body length to dry mass following the length-weight relationships (McCauley, 1984), and applied a factor of 0.4 to convert dry mass to carbon mass (Reiss & Schmid-Araya, 2008).

Due to the high time demand for taxonomic identification, counting and size measurements, 3 randomly selected replicates per all treatment combinations were processed. These data (i.e., N = 3 per treatments) were then used in the analyses of species richness and community composition. For zooplankton, the dominant taxa (i.e., cladoceran genera, Cyclopoida, and Calanoida) were identified in all 5 replicates per treatments, which we used to calculate taxa-specific biomass and to analyse resistance and recovery of zooplankton. As rotifers occurred in very low abundances from t_2_ till the end of the experiment, we considered the biomass of crustaceans as a representative proxy for the total zooplankton biomass.

### 2.4. Data analysis

To illustrate temporal dynamics of daily mean temperatures and Chla-a fluorescence (proxy for phytoplankton biomass) over the experimental duration, we fitted smoothed conditional mean curves based on generalised additive models (GAM) with the *stat_smooth* function (using mgcv gam fitting and formula: y∼s(x,k=12)) of the ‘ggplot2’ R package (Wickham et al., 2021). To visualise the immediate effect of the dispersal treatments (at t_1_, before the heatwave manipulation) on phytoplankton and zooplankton alpha diversity, we compared rarefied (i.e., testing for density-independent differences in taxon richness) mean taxon richness between the treatments with and without dispersal manipulations by the ‘mobr’ R package (McGlinn et al., 2021). Effect sizes were calculated as the mean absolute differences between treatments 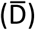, and p-values were determined by a Monte Carlo permutation procedure (n = 1,000 permutations) following the framework described in McGlinn et al. (2019). To visualise the effect of connectivity on gamma diversities, we constructed sample-based rarefaction curves for both phyto- and zooplankton based on 1,000 random permutations using the *specaccum* function of the ‘vegan’ R package (Oksanen et al., 2020).

To assess the effect of heatwave (H+) and dispersal (M+, Z+) on the resistance and recovery of plankton biomass, we fitted linear models (LMs). Resistance was quantified as the biomass change (i.e., difference) between the time points before (t_1_) and at the end (t_2_) of the heatwave (i.e., the response to the heatwave). Recovery after the heat wave was estimated as the change in biomass between the two time points in the recovery phase (t_3_ and t_4_), where total plankton biomass in the H+ treatments started to increase after minimum values around t_3_ (Figure S3). For both resistance and recovery, we created separate LMs with the following dependent variables: (1) change of total plankton biomass, i.e., the sum of zooplankton carbon mass and POC (including all microorganisms <100 μm), (2) change of zooplankton biomass (carbon mass), (3) change of Chl-*a*, and (4) change of POC. In all models, heatwave (two levels: H+, H-) and connectivity (four levels: M-Z-, M+Z-, M-Z+, and M+Z+), as well as their two and three-way interactions were included as fixed factors. To account for any potential differences in initial biomass values, mean-centred initial biomass (i.e., t_1_ for resistance and t_3_ for resilience) was also included in all models as a predictor. Assumptions of normality and homoscedasticity of residuals were assessed by diagnostic plots.

POC is considered a proxy for microbial biomass, i.e., the sum of bacteria, protists and phytoplankton (Davidson et al., 2002; Gerhard et al., 2022; Poxleitner et al., 2016). Accordingly, we used POC (<100 μm fraction) as a proxy for the biomass of microorganisms. Even though POC may contain some detritus and even smaller zooplankton such as rotifers, we still consider it as a representative proxy in our experiment for the following reasons. First, as we were primarily interested in the planktonic communities, we did not include sediment in the mesocosms, which could have been resuspended and contribute to POC substantially. Second, given the enclosed nature of the mesocosms, negligible amounts of detritus input from the surrounding terrestrial habitats could be expected. Third, small zooplankton, such as rotifers, were rare from t_2_ onwards, therefore, we considered the contribution of the zooplankton size category between 30 and 100 μm to POC as negligible. Additionally, POC generally corresponded well to phytoplankton carbon mass as indicated by significant positive correlations at the focal sampling dates t_1_ and t_4_ (at t_5_, algal biomasses were generally very low, hence the relationship was also weaker; Figure S4).

We also tested the responses of the daily Chl-*a* fluorescence (as dependent variable) to the experimental treatments to analyse resistance and recovery of phytoplankton biomass with a higher temporal resolution. To this end, three separate linear mixed-effect models (LMEMs) were constructed covering the experimental period from the start of the experimental heatwave until the end of the experiment. To determine resistance, the first model was fitted to the data of the heatwave period (from day 10 to day 21, n = 12 days). As microorganisms such as phytoplankton can rapidly respond to disturbances due to high population growth rates, we decided to analyse short-term and delayed effects of the experimental treatments by building separate models for the first and second part of the recovery phase. We tested short-term recovery immediately after the culmination phase of the heatwave (from day 22 to day 33, n = 12 days). We chose this period as Chl-*a* fluorescence values were lowest at the end of the experimental heatwave in all treatment groups, and started to increase from day 22 (Figure S2). In the third model, we analysed the temporal pattern of Chl-*a* in the second part of the recovery phase (from day 34 to day 44, n = 11 days) to test for any lagged effects of the experimental treatments. In all LMEMs, we included the experimental treatments, time (i.e., day centred on the minimum value of the given period), as well as their two and three-way interactions as fixed factors. Here, a significant interaction with time can be interpreted as a temporal trend within the experimental treatments. We also built in zooplankton biomass observed at the beginning at each of the three analysed periods to test for the effect of top-down control. We set individual mesocosm as random intercept and included the AR(1) error structure to account for temporal autocorrelation (Pinheiro & Bates, 2000). Model comparison by the Akaike Information Criterion indicated that accounting for temporal autocorrelation improved the model fit in all models. Chl-a fluorescence data were log-transformed in all models to normalise residuals and improve homoscedasticity of variances. Linear mixed-effects models were constructed with the *lme* function of the ‘nlme’ R package (Pinheiro et al., 2022). Marginal and conditional R^2^ of the models were calculated by the *r*.*squaredGLMM* function of the ‘MuMIn’ R package (Bartoń, 2022).

To reveal how food web structure and trophic transfer efficiency changed in response to the experimental manipulation, we tested treatment-specific differences in the ratio of zooplankton carbon mass to POC (i.e., carbon mass of microorganisms). This biomass ratio between organisms with higher and lower trophic positions indicates whether the food webs are more top- or bottom-heavy (Shurin et al., 2012). We constructed separate LMs for all four time points with heatwave and dispersal as fixed factors, treated the same way as in the models for resistance and recovery. We assured that normality and homoscedasticity of residuals were met by using diagnostic plots.

In order to test which taxa may be responsible for the observed treatment effects, we analysed the changes in community composition (based on taxon-specific biomasses) and tested for significant associations between specific taxa and treatments. We first visualised phytoplankton and zooplankton composition across all treatment combinations with non-metric multidimensional scaling (NMDS) based on Bray-Curtis dissimilarity matrices. We then tested for significant treatment effects using permutational multivariate analysis of variance (PERMANOVA, Anderson, 2001), based on Bray-Curtis distances and 1,000 random permutations. To identify taxa with the strongest contribution to compositional changes among significantly different treatments, we performed a similarity percentages (SIMPER) analysis. Significant associations between taxa and treatments were tested by 1000 random permutations. The analyses were performed with the functions *metaMDS* (NMDS), *adonis2* (PERMANOVA), and *simper* in the ‘vegan’ package. All data analyses and visualisations were performed in R version 4.0.2 (R Core Team, 2020).

## 3. Results

During the 7-days long culmination of the heatwave period, we maintained an approximately 5°C offset between the H+ and H-treatments (Figure 1), which resulted in mean water temperatures of 25.5 ± 1.3 °C (mean ± SD, N = 20) in the H+ treatment and 20.9 ± 1.2 °C (mean ± SD, N = 20) in the H-treatment. Experimental dispersal (M+) had an immediate positive effect on phytoplankton taxonomic richness expressed as both alpha and gamma diversities (Figure 2a-b). Rarefied richness at the local scale (i.e., alpha-diversity) was significantly higher compared to the control (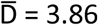, p < 0.05; Figure 2a). Dispersal (Z+) did not result in increased zooplankton alpha-diversity relative to the control (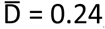, p = 0.62; Figure 2c), even though there were overall more zooplankton species (i.e., greater gamma-diversity) in the Z+ treatments (Figure 2d).

**Figure 2.**
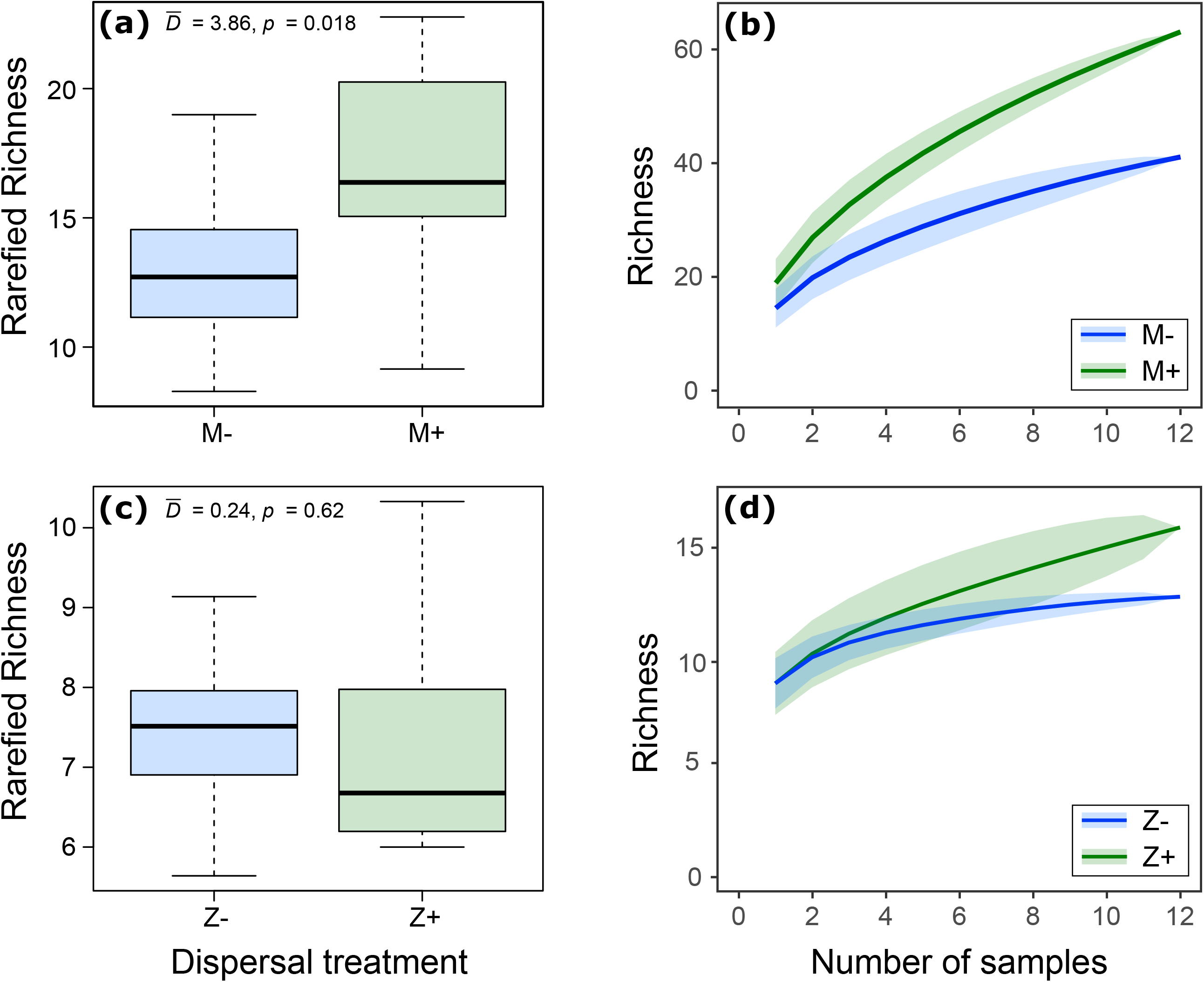
Richness of (**a-b**) phytoplankton and (**c-d**) zooplankton at time point t_1_, i.e., after applying the dispersal treatments (M+ and Z+), but prior to the experimental heatwave. (a) Boxplots illustrate that rarefied richness of phytoplankton at the local scale (i.e., alpha-diversity) increased significantly in the presence of dispersal (M+). 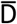 stands for mean absolute differences between treatments, the *p*-values are obtained from a Monte Carlo permutation procedure with 1000 permutations. (**b**) Species accumulation curves (mean ± SD based on 1000 random permutations) show that M+ also increased gamma-diversity. (**c**) Boxplots illustrate that rarefied richness of zooplankton at the local scale (i.e., alpha-diversity) was not significantly different among the dispersal treatments. (**d**) Species accumulation curves (mean ± SD based on 1000 random permutations), however, show a higher zooplankton gamma-diversity with Z+.

The H+ treatment suppressed total plankton biomass (LM, p < 0.001, Table 1, Figure 3, Figure S3). This significant decrease in biomass was primarily driven by zooplankton (LM, p < 0.001, Table 1). Especially the biomass of cladocerans declined strongly in response to H+ (LM, p < 0.001), while the negative heatwave effect was weaker on copepods (LM, p = 0.07, Table S3, Figure S5). Besides the biomass decline, the ratio of egg-carrying females in the populations of dominant cladocerans, *D*. cf. *longispina*, also dropped from 35.1 ± 9.4 (t_2_) to 0.0 ± 0.0% (t_5_) in the H+ treatments (mean ± SD, N = 20), but this was similar in the control treatments (mean ± SD at t_2_: 31.6 ± 8.0%, t_5_: 0.3 ± 0.6%, N = 20). In contrast to the negative effects of H+ on total plankton and zooplankton biomasses, we did not find any significant effect on Chl-*a* concentration and POC (LM, p > 0.05, Table 1), nor on the pattern of daily Chl-*a* fluorescence (LMEM, p>0.05, Table 2). Our results did not provide evidence for enhanced resistance to H+ following dispersal from the regional species pool. None of the experimental dispersal treatments mitigated the heatwave-driven decline observed in total plankton biomass and zooplankton (LM, p > 0.1, Table 1, Figure 3). Intriguingly, zooplankton dispersal (M-Z+ and M+Z+) had a negative effect on zooplankton resistance, i.e., suppressed biomass during the heatwave manipulation (LM, p < 0.05, Table 1).

**Table 1.** Summary statistics (estimated parameters, standard error and p values) of multiple linear regression models testing the effect of the experimental treatments (H+: heatwave, M+: dispersal of microorganisms, Z+: dispersal of zooplankton) on resistance and recovery of plankton community biomass. Resistance is defined as the change in biomass during the heatwave (i.e., difference between t_1_ and t_2_), while recovery as the change in the post-heatwave period (i.e., difference between t_3_ and t_4_). Abbreviations: Chl-*a* – chlorophyll *a* concentration, POC – particulate organic carbon, initial – mean-centred initial biomass. The models for Chl-*a* and the model testing recovery of zooplankton biomass are based on log-transformed data. Significant results (p < 0.05) are highlighted in bold, while marginally significant ones (p < 0.1) with italics.

| | Total plankton biomass | | | Zooplankton biomass | | | log Chl-$a$ | | | POC | | |
| --- | --- | --- | --- | --- | --- | --- | --- | --- | --- | --- | --- | --- |
| | est. | SE | $p$ | est. | SE | $p$ | est. | SE | $p$ | est. | SE | $p$ |
| **Resistance** |  |  |  |  |  |  |  |  |  |  |  |  |
| intercept | -7.92 | 7.48 | 0.298 | **15.40** | **3.45** | **<0.001** | -0.38 | 0.27 | 0.168 | **-23.05** | **5.94** | **<0.001** |
| initial | **-1.03** | **0.15** | **<0.001** | **-1.27** | **0.13** | **<0.001** | **-0.45** | **0.06** | **<0.001** | **-0.86** | **0.14** | **<0.001** |
| H+ | **-26.23** | **10.67** | **0.020** | **-24.88** | **4.89** | **<0.001** | -0.33 | 0.35 | 0.357 | -1.91 | 1.07 | 0.822 |
| M+Z- | -3.54 | 10.60 | 0.741 | -7.84 | 4.97 | 0.125 | -0.40 | 0.35 | 0.272 | 2.55 | 1.08 | 0.767 |
| M-Z+ | -6.35 | 10.59 | 0.553 | **-11.06** | **4.96** | **0.033** | -0.26 | 0.35 | 0.465 | 2.96 | 1.07 | 0.727 |
| M+Z+ | -10.30 | 10.58 | 0.338 | **-13.01** | **4.91** | **0.013** | -0.19 | 0.36 | 0.585 | 4.34 | 1.18 | 0.611 |
| H+ × M+Z- | 15.71 | 14.99 | 0.303 | 6.36 | 7.00 | 0.370 | 0.78 | 0.50 | 0.132 | 13.11 | 1.56 | 0.284 |
| H+ × M-Z+ | -0.88 | 15.03 | 0.954 | 6.33 | 6.91 | 0.367 | 0.46 | 0.50 | 0.361 | -4.91 | 1.52 | 0.685 |
| H+ × M+Z+ | 5.25 | 14.96 | 0.728 | 9.41 | 6.97 | 0.187 | 0.67 | 0.50 | 0.189 | -6.40 | 1.63 | 0.595 |
| **R<sup>2</sup>** |  | 0.62 |  |  | 0.83 |  |  | 0.57 |  |  | 0.53 |  |
| **Recovery** |  |  |  |  |  |  |  |  |  |  |  |  |
| intercept | 4.07 | 9.87 | 0.682 | **0.48** | **0.18** | **<0.01** | -0.04 | 0.22 | 0.851 | -5.96 | 8.11 | 0.468 |
| initial | **-0.78** | **0.25** | **<0.01** | **-0.62** | **0.18** | **<0.01** | **-0.76** | **0.13** | **<0.001** | **-0.95** | **0.21** | **<0.001** |
| H+ | 3.26 | 14.3 | 0.821 | -0.10 | 0.26 | 0.715 | 0.51 | 0.32 | 0.120 | 8.30 | 11.5 | 0.477 |
| M+Z- | 8.67 | 13.46 | 0.524 | -0.00 | 0.22 | 0.985 | 0.37 | 0.31 | 0.244 | 8.00 | 11.5 | 0.490 |
| M-Z+ | 1.86 | 13.49 | 0.891 | 0.25 | 0.23 | 0.267 | -0.01 | 0.31 | 0.976 | -5.92 | 11.4 | 0.477 |
| M+Z+ | 19.30 | 13.81 | 0.172 | 0.04 | 0.23 | 0.866 | 0.26 | 0.31 | 0.401 | 16.77 | 11.6 | 0.158 |
| H+ × M+Z- | -1.34 | 19.19 | 0.945 | -0.08 | 0.33 | 0.82 | -0.15 | 0.46 | 0.525 | 7.14 | 16.2 | 0.670 |
| H+ × M-Z+ | 8.81 | 19.24 | 0.650 | -0.43 | 0.32 | 0.188 | 0.29 | 0.44 | 0.732 | 18.84 | 16.6 | 0.255 |
| H+ × M+Z+ | -10.17 | 19.24 | 0.600 | -0.1 | 0.32 | 0.74 | -0.20 | 0.44 | 0.669 | -7.83 | 16.2 | 0.632 |
| **R<sup>2</sup>** |  | 0.29 |  |  | 0.28 |  |  | 0.55 |  |  | 0.48 |  |

**Table 2.** Summary statistics (estimated parameters, standard error and p values) of linear mixed-effects models testing the effects of the experimental treatments (H+: heatwave, M+: dispersal of microorganisms, Z+: dispersal of zooplankton) on resistance and recovery of phytoplankton biomass (measured as log-transformed chlorophyll *a* in vivo fluorescence). Models were built for the period of the experimental heatwave (n = 12 days between days 10–21 and t_2_, ‘Resistance’), and for the first (n = 12 days between days 22–33, ‘Recovery 1’) and second part (n = 11 days between days 34–44, ‘Recovery 2’) of the recovery phase. Significant results (p < 0.05) are highlighted in bold, while marginally significant ones (p < 0.1) with italics.

|  | Resistance |  |  | Recovery 1 |  |  | Recovery 2 |  |  |
| --- | --- | --- | --- | --- | --- | --- | --- | --- | --- |
|  | est. | SE | *p* | est. | SE | *p* | est. | SE | *p* |
| intercept | **4.07** | 0.24 | **<0.001** | **2.73** | 0.64 | **<0.001** | 4.68 | 0.49 | **<0.001** |
| H+ | 0.22 | 0.27 | 0.431 | -0.16 | 0.54 | 0.766 | -0.79 | 0.52 | 0.143 |
| M+Z- | 0.23 | 0.27 | 0.410 | 0.53 | 0.46 | 0.259 | 0.30 | 0.50 | 0.556 |
| M-Z+ | 0.10 | 0.27 | 0.724 | *0.80* | *0.46* | *0.097* | -0.48 | 0.49 | 0.341 |
| M+Z+ | -0.13 | 0.27 | 0.629 | 0.31 | 0.48 | 0.532 | 0.40 | 0.49 | 0.810 |
| day | **-0.19** | 0.03 | **<0.001** | **0.20** | 0.04 | **<0.001** | **-0.09** | 0.04 | **0.022** |
| ZPB | **-0.01** | -0.01 | **<0.01** | *-0.01* | *0.00* | *0.087* | **-0.02** | 0.01 | **0.027** |
| H+ × M+Z- | -0.28 | 0.38 | 0.474 | -0.70 | 0.65 | 0.290 | 0.44 | 0.71 | 0.543 |
| H+ × M-Z+ | -0.26 | 0.38 | 0.504 | -0.26 | 0.65 | 0.696 | 0.48 | 0.70 | 0.494 |
| H+ × M+Z+ | -0.15 | 0.38 | 0.699 | -0.44 | 0.66 | 0.512 | 0.59 | 0.70 | 0.401 |
| day × H+ | 0.01 | 0.04 | 0.822 | -0.08 | 0.06 | 0.181 | **0.13** | 0.06 | **0.024** |
| day × M+Z- | -0.01 | 0.04 | 0.837 | -0.09 | 0.06 | 0.151 | -0.03 | 0.06 | 0.546 |
| day × M-Z+ | 0.02 | 0.04 | 0.663 | *-0.12* | *0.06* | *0.054* | 0.03 | 0.06 | 0.658 |
| day × M+Z+ | 0.06 | 0.04 | 0.120 | -0.05 | 0.06 | 0.375 | -0.03 | 0.06 | 0.649 |
| day × H+ × M+Z- | 0.00 | 0.06 | 0.936 | **0.18** | **0.08** | **0.030** | -0.01 | 0.08 | 0.917 |
| day × H+ × M-Z+ | 0.02 | 0.06 | 0.759 | 0.08 | 0.08 | 0.347 | -0.04 | 0.08 | 0.583 |
| day × H+ × M+Z+ | -0.01 | 0.06 | 0.806 | 0.13 | 0.08 | 0.126 | -0.11 | 0.08 | 0.179 |
| Marginal R<sup>2</sup> |  | 0.57 |  |  | 0.32 |  |  | 0.33 |  |
| Conditional R<sup>2</sup> |  | 0.57 |  |  | 0.74 |  |  | 0.33 |  |

**Figure 3.**
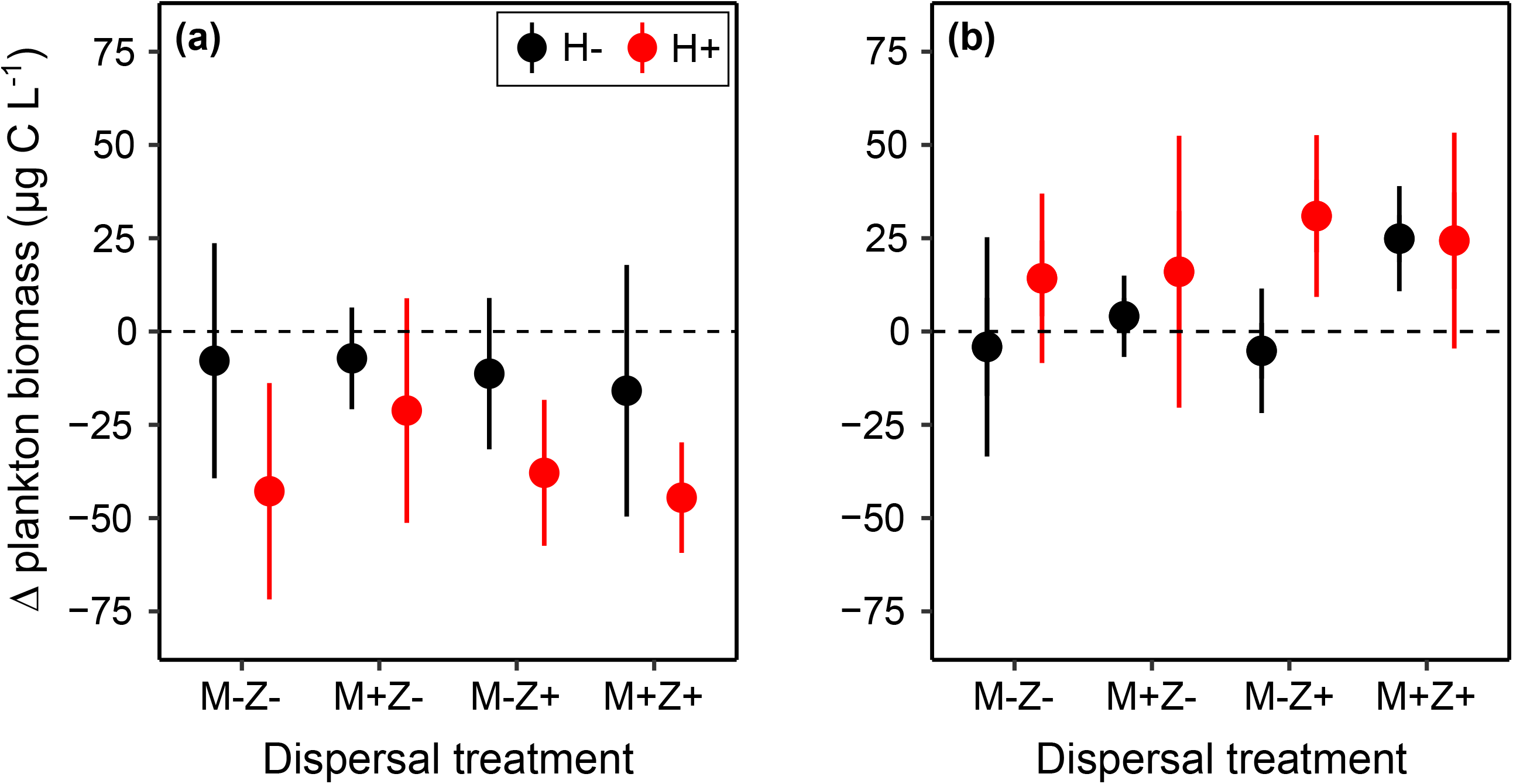
Resistance and recovery of the plankton community measured as the change of total plankton biomass (expressed in carbon mass; mean ± SD) (**a**) during the heatwave (i.e., difference between t_1_ and t_2_) and (**b**) the recovery period (difference between t_3_ and t_4_). N = 5 for all treatment combinations. (**a**) The experimental heatwave (H+) had a significant effect on the change of total plankton biomass (p < 0.001 and p = 0.02, Table 1), by increasing the degree of biomass decrease relative to the control (H-) conditions. The dispersal treatments (microorganism: M- and M+; zooplankton: Z- and Z+) did not influence the changes of biomass. (**b**) None of the experimental treatments had a significant influence on the changes of total plankton biomass in the recovery phase. Summary statistics are presented in Table 1.

Although there was no evidence for community resistance to heatwave, we found that microorganism dispersal contributed to a faster growth of phytoplankton biomass (measured as Chl-*a* fluorescence) in the presence of heatwave. This was particularly visible in the first part of the post-heatwave recovery phase, indicated by the significant interaction between day × H+ × M+Z- (LMEM, p<0.05, Table 2, Figure 4). The insurance effect of microbial dispersal was no longer detectable in the second part of the recovery phase, but H+ enhanced phytoplankton biomass towards the end of the experiment (i.e., significant day × H+ interaction, LMEM, p<0.05, Table 2, Figure S6). Moreover, zooplankton biomass had a significant negative effect on Chl-*a* fluorescence (i.e., negative effect on the intercept, LMEM, p<0.05, Table 2), indicating that top-down control was a major driver of phytoplankton dynamics in the recovery phase. There was no effect of dispersal on the recovery of total plankton biomass, zooplankton, and the microorganism biomass proxies (Chl-*a*, POC) between t_3_ and t_4_ in the recovery phase (LM, p>0.05, Table 1).

**Figure 4.**
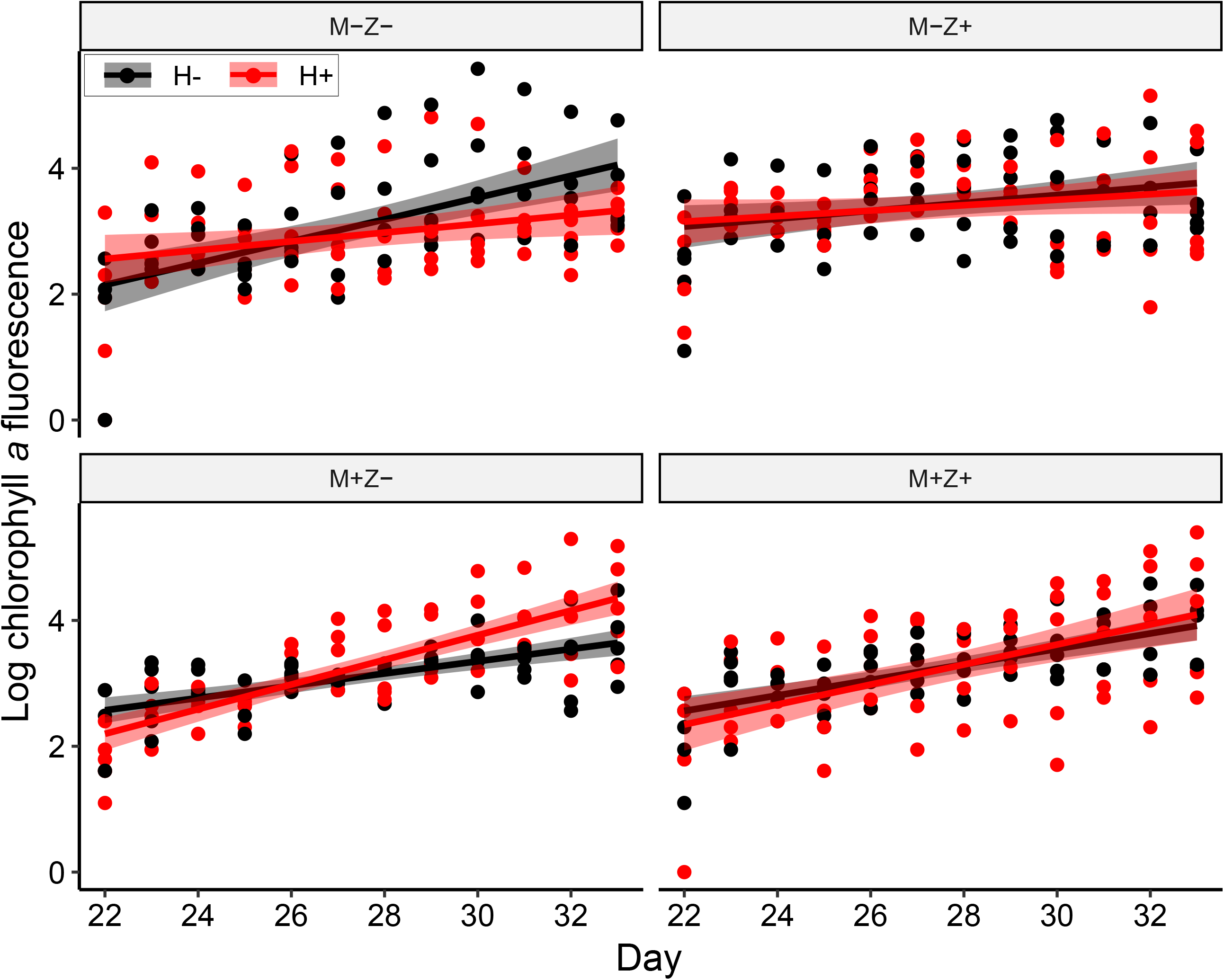
Increases in daily in vivo fluorescence of chlorophyll a (proxy for phytoplankton biomass) following the experimental heatwave, grouped according to the dispersal treatments (microorganism: M- and M+; zooplankton: Z- and Z+) and coloured according to the heatwave treatment (H- and H+). N = 5 for all treatment combinations per day. Solid lines represent fitted linear models (error bands: 95% confidence intervals) to visualise temporal trends. Based on linear mixed-effects models, recovery in the H+ treatments after the heatwave was enhanced by M+ (LMEM: significant day × H+ × M+Z-, p < 0.05, Table 2).

Planktonic food webs became more bottom-heavy as a response to H+, indicated by the significant negative effect on zooplankton carbon mass to POC ratios (LM, p = 0.007, Figure 5, Table S4). This effect became evident after the heat wave and lasted until the end of the experiment (Table S4).

**Figure 5.**
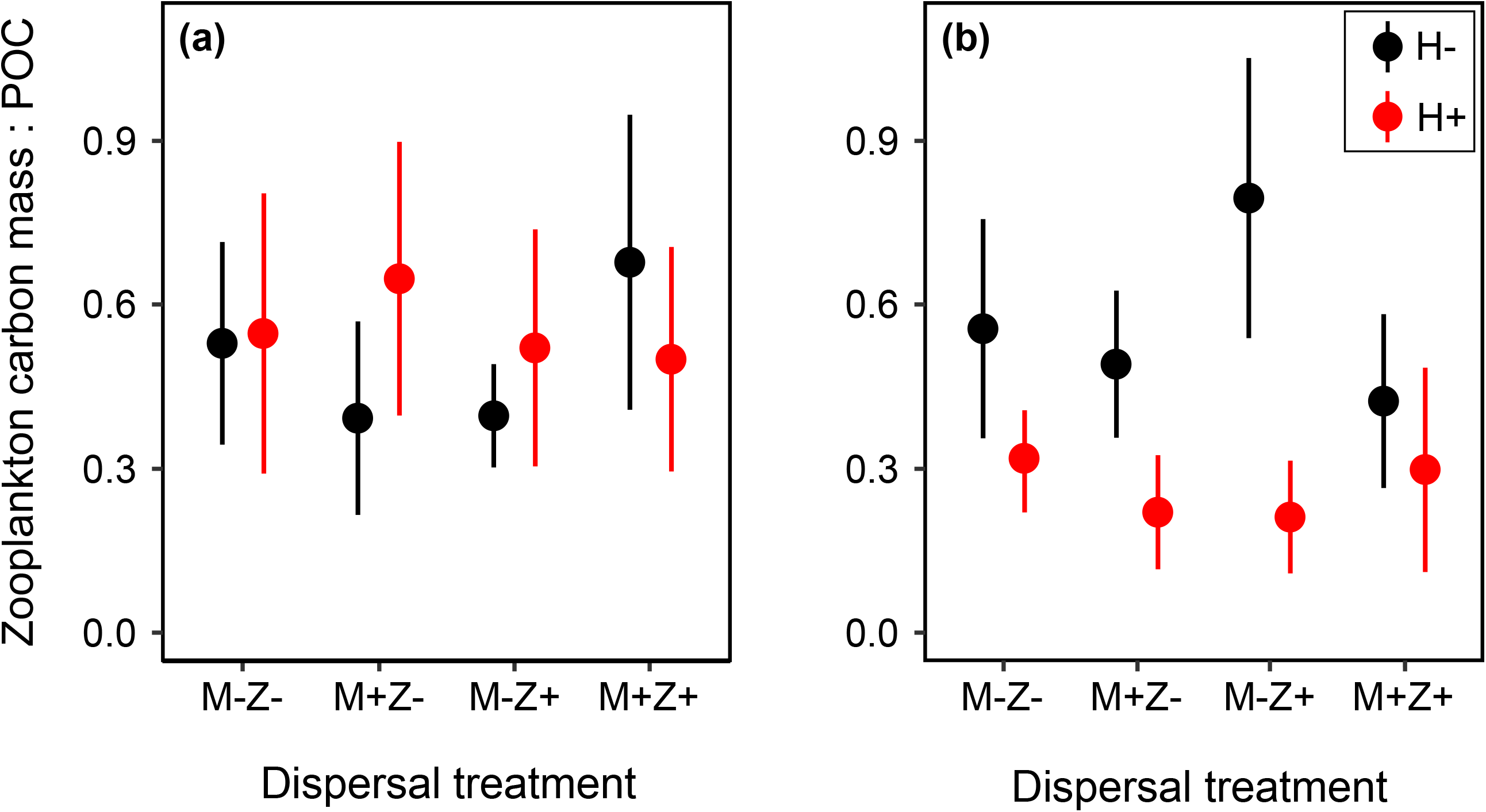
(**a**) Food web structure at the beginning (sampling point t_1_) and (**b**) the end (t_4_) of the experiment expressed as the ratio (mean ± SD) of carbon mass between zooplankton and POC (proxy for microorganisms). N = 5 for all treatment combinations. While the experimental treatments had no significant effect at t_1_, the ratio decreased as a response to the heatwave (H+) at t_4_, i.e., food webs shifted from top-heavy to more bottom-heavy in the H+ treatments (LM with log-transformed data, p < 0.007, see Table S4 for the statistical summary).

In total, 137 phytoplankton taxa were identified. Based on their mean biomasses at the focal sampling dates, the communities were dominated by the diatom *Nitzschia* sp., the chrysophyte *Chromulina* sp., and the green algae *Scenedesmus* group *Acutodesmus*, an unidentified single-celled green alga (diameter: 2.5 μm), and *Mougeotia* sp. (Table S5). We found 20 zooplankton taxa (Table S2), of which four were dominant (based on frequency of occurrence and contribution to total biomass) during the experiment. These taxa were the cladocerans *Daphnia* cf. *longispina* and *Bosmina longispina*, and the copepods *Eudiaptomus gracilis* and *Cyclops abyssorum*. The most dominant taxon was *Daphnia* cf. *longispina*, which accounted for 47.5 ± 18.9 (at t_2_) to 61.5 ± 15.1% (at t_5_) of total zooplankton biomass over the experiment (mean ± SD, N = 24).

The heatwave (H+) and microbial dispersal (M+) treatments both had a significant effect on community composition of phytoplankton, which became evident towards the end (t_4_) of the experiment (PERMANOVA, p < 0.05, Figure 6, Table 3). Among the most influential taxa, in terms of total explained variation across treatments, green algae belonging to *Scenedesmus* group *Acutodesmus* and Ulotrichales were positively associated with H+ (SIMPER, p < 0.05, Table S6). Another green alga, *Mougeotia*, exhibited significantly lower biomass values in the M-vs M+ treatment (SIMPER: p < 0.05, Table S6). Zooplankton community composition was only affected by H+, which was evident already from t_2_ (PERMANOVA: p < 0.01, Table 3, Figure S7) and lasted until t_4_ (PERMANOVA, p < 0.01, Figure 6, Table 3). At both t_2_ and t_4_, *Daphnia* cf. *longispina* had the highest contribution to the overall dissimilarity across treatments, while at the same time, it had significantly lower biomasses in the H+ treatment (SIMPER, p < 0.01, Table S6). Another cladoceran, *Bosmina longispina*, was negatively associated with H+ (PERMANOVA, p < 0.01, Table S6).

**Figure 6.**
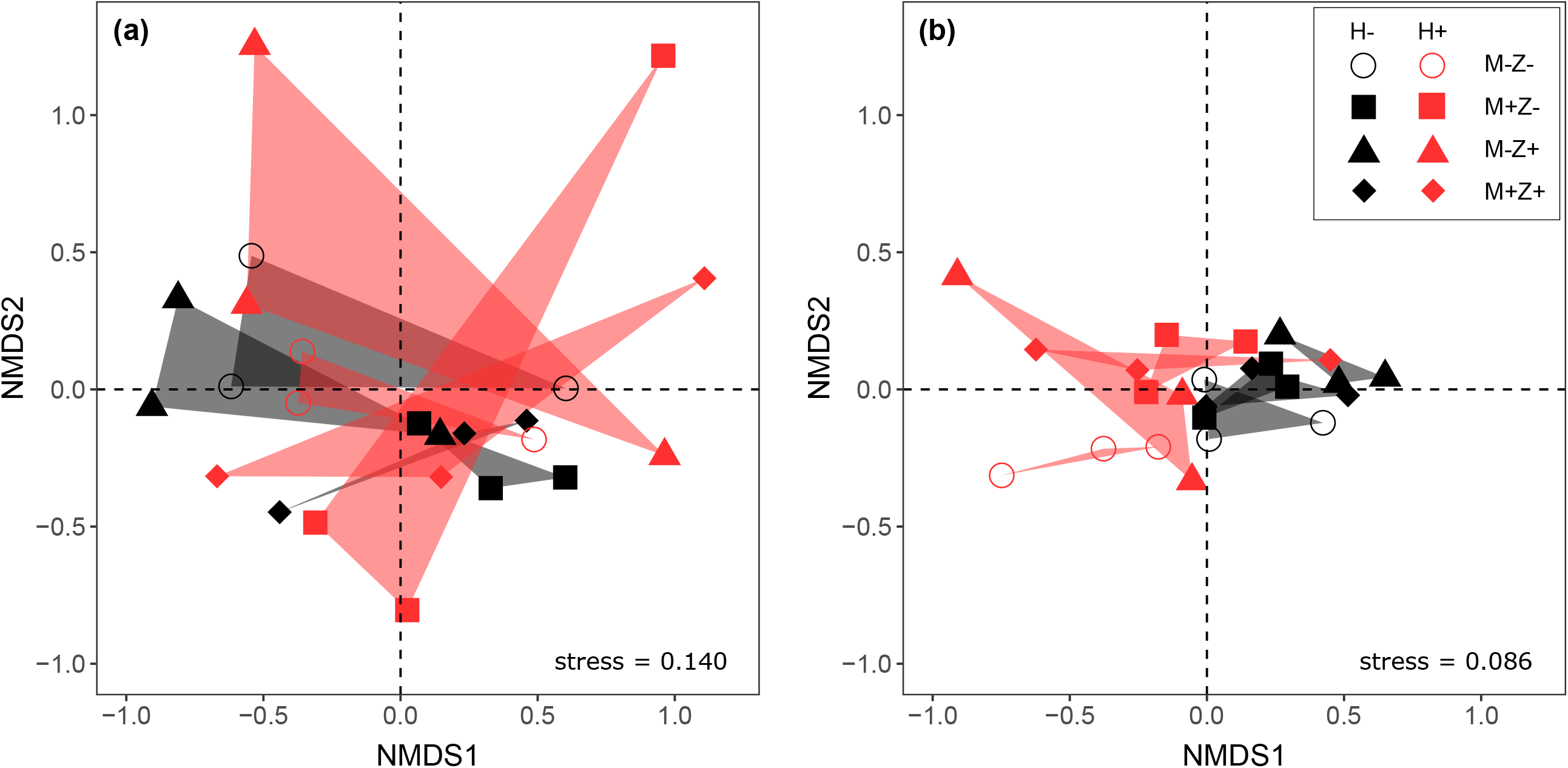
NMDS plots illustrating the effects of heatwave (H-, H+) and dispersal treatments (microorganisms: M-, M+, zooplankton: Z-, Z+) on (**a**) phytoplankton and (**b**) zooplankton community composition (based on biomasses) towards the end of the experiment (t_4_). N = 3 for all treatment combinations. Phytoplankton community composition was significantly influenced by both H+ (PERMANOVA: p < 0.05) and M+ (p < 0.05), while zooplankton by H+ only (p < 0.01). Results of PERMANOVAs are presented in Table 2.

**Table 3.**
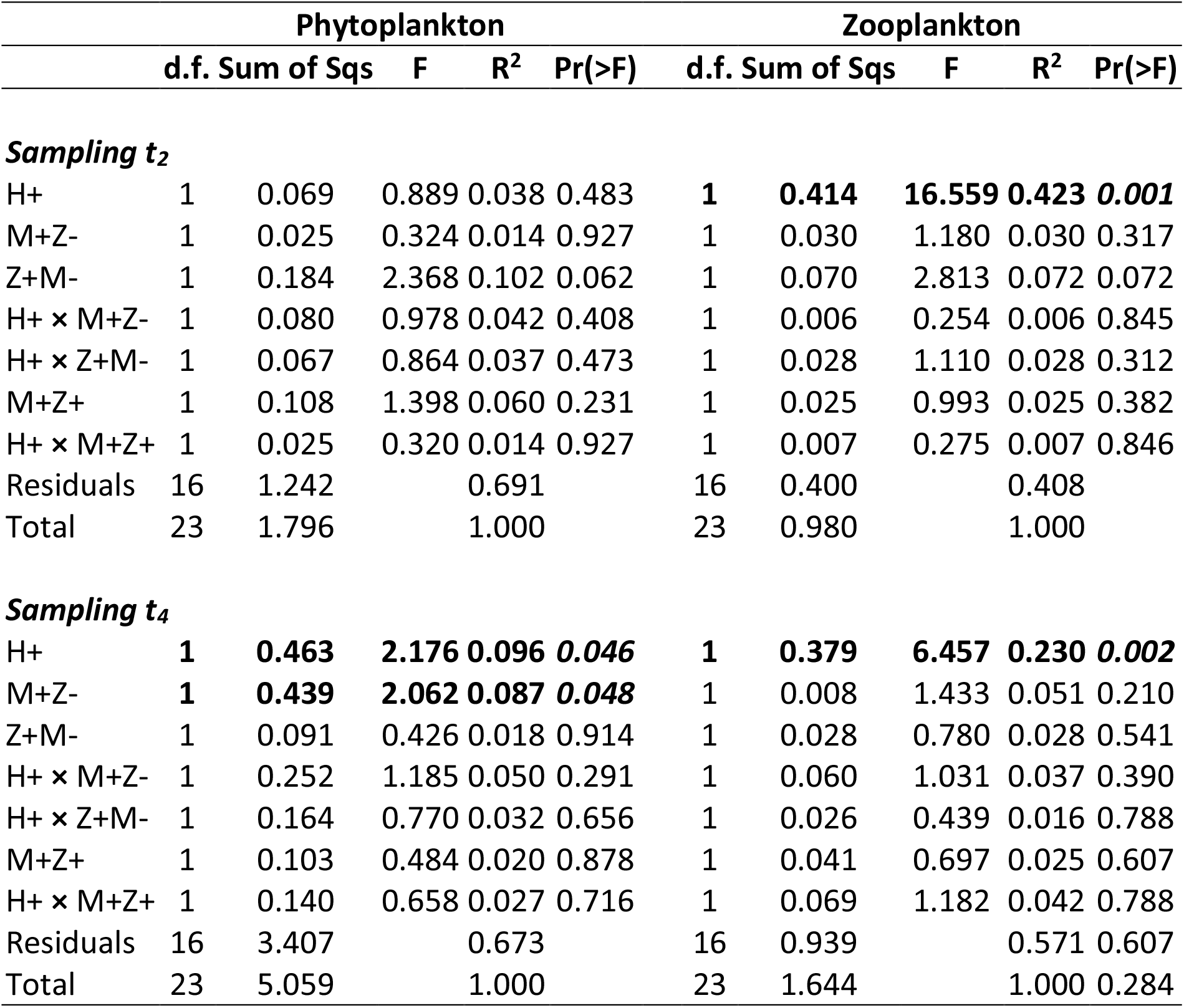
Summary statistics of PERMANOVA testing for treatment-specific differences in (biomass based) community composition of phytoplankton and zooplankton at the end of the heatwave (t_2_) and towards the end of the experiment (t_4_). Abbreviations of treatments: H+: heatwave, M+: microorganism dispersal, Z+: zooplankton dispersal. Significant (p < 0.05) results are highlighted in bold.

## 4. Discussion

The experimental heatwave had a strong negative effect on total community biomass supporting our first hypothesis. This negative effect was primarily driven by the decline in zooplankton biomass. The heatwave-driven disruption of the trophic structure resulted in a shift from top-heavy to more bottom-heavy food webs. Altogether, these findings are in agreement with our second hypothesis. Weakened top-down control contributed to an elevated phytoplankton biomass, however, this effect became visible only about two weeks following the heatwave. These results illustrate that heatwave impacts may only amplify over time as a result of cascading interactions (Ross et al., 2022).

Experimental dispersal did not have an apparent effect on biomass resistance of microbial and zooplankton communities. Towards the end of the experimental period, total plankton biomass recovered gradually by reaching similar levels as in the control treatment, indicating high capacity for community resilience. In contrast to our third hypothesis, there was no evidence for the buffering effect of dispersal on zooplankton recovery. However, microbial dispersal had a positive effect on the recovery of biomass at the level phytoplankton following the heatwave. This can be interpreted as spatial insurance, as higher biomass of primary producers may provide a basis for faster recovery of secondary producers and hence total community biomass over time. Altogether, these results suggest that dispersal can buffer the negative effect of heatwaves, and its effect depends on the trophic level of organisms. Spatial insurance may be more evident in unicellular organisms than in larger organisms at upper trophic levels (Limberger et al., 2019).

The decline of total plankton biomass during the heatwave is consistent with the prediction that increasing temperatures reduce total community biomass (O’Connor et al., 2009). This prediction is based on the differential temperature scaling of respiration- and photosynthesis-limited metabolism, which implies a greater sensitivity and therefore stronger responses of heterotrophic organisms to temperature changes compared to autotrophs (López-Urrutia et al., 2006; O’Connor et al., 2011). Increased grazing pressure with warming, driven by increased metabolic demands, generally results in a shift towards top-heavy food webs (O’Connor et al., 2009; Shurin et al., 2012; Velthuis et al., 2017). However, a greater proportion of consumers is expected to result in a decline of total biomass given the inefficient conversion of phytoplankton to consumer biomass (Persson et al., 2007; Slobodkin, 1959). In contrast, we found that food webs became more bottom-heavy (resource-controlled) following the experimental heatwave. Our results therefore suggest that temperature either had a direct negative impact on secondary producers, or that resource limitation intensified to an extent which resulted in an abrupt decline in their biomass.

The biomass of crustacean zooplankton, especially of cladocerans (dominated by *Daphnia* cf. *longispina*) declined strongly as a result of the heatwave, which was the major driver of the observed decline of total plankton biomass. While cladocerans can rapidly respond to temperature stress by adjusting their physiology (e.g., Yampolsky et al., 2014), the abrupt decline in biomass may indicate a failed acclimation. Besides direct temperature effects, the timing and magnitude of temperature fluctuations, and their interactions with food-limited periods are crucial for zooplankton phenology (Huber et al., 2010). In our study, the experimental heatwave coincided with a ‘clear-water phase’ (in all treatments), i.e., a trophic cascade where zooplankton increased to the point it became strongly food-limited (Lampert et al., 1986). This was evidenced by the fact that the lowest phytoplankton biomasses (Figure S2) and, at the same time, the highest zooplankton biomasses in the control (i.e., H-) treatments (Figure S5) were observed at the end of the experimental heatwave period. This suggests that the effect of elevated temperature was amplified by starvation of key consumers and that causal relationships may, at least partly, be based on indirect effects of resource shortage. The lack of egg carrying females even in the control (i.e., H-) mesocosms further indicated that *Daphnia* did not have sufficient food supply to invest into reproduction. While the experimental heatwave clearly affected population dynamics of cladocerans, it had a weaker impact on copepod biomass (marginally significant negative effect). This may be related to the fact that copepods have a broader dietary niche which includes rotifers and ciliates (Adrian & Schneider-Olt, 1999; Brandl, 2005). Besides, copepods are generally more buffered against starvation due to their better ability to accumulate high amounts of storage lipids (Brett et al., 2009; Lampert & Muck, 1985). Taken together, these findings also indicate that copepods may be generally more robust to temperature fluctuations than cladocerans.

Zooplankton dispersal did not have a clear effect on zooplankton community composition, and did not provide a buffering effect on zooplankton biomass. Instead of an expected enhanced resistance to the heatwave, there was an intriguing negative effect of zooplankton dispersal on zooplankton biomass. Metacommunity theory predicts that excess dispersal can reduce local community biomass by maintaining the presence and dominance of low-performing species in local habitats over better-suited species (Leibold et al., 2017; Matthiessen & Hillebrand, 2006). However, the single dispersal event with a low density of introduced zooplankters represented a rather low dispersal rate to considerably modify density-dependent processes. Alternative mechanisms might have negatively influenced the crustacean biomass in our Z+ treatments, for example, introduction of parasites along with the regional dispersal inoculum.

In contrast to zooplankton dispersal, microbial dispersal enhanced growth of phytoplankton biomass (i.e., Chl-*a* fluorescence) following the heatwave. It also had a significant effect on phytoplankton community composition and a much stronger effect on phytoplankton taxon richness than zooplankton dispersal. A possible explanation for the differential effects of zooplankton and microbial dispersal may be related to the timing of dispersal. Colonisation success depends on arriving at a window of opportunity, i.e., empty niche space in the resident community (Clark & Johnston, 2011; Symons & Arnott, 2014; Thompson & Gonzalez, 2017). Generally, limited resource use efficiency (i.e., high amount of available resources) in a local community increases the opportunity for invasion (Davis et al., 2000). Though dispersal timing in our experiment coincided for phytoplankton and zooplankton, local phytoplankton and zooplankton communities differed in their successional stages, creating different windows of opportunity for invasion. By the time of dispersal, plankton communities were strongly consumer controlled, and high zooplankton grazing depleted food resources during the period of the heatwave (i.e., even in the H-treatments). Therefore, zooplankters introduced from the regional pool were facing an environment with intense competition for limited resources. It is likely that the probability for successful establishment would have been higher following the heatwave, when monopolization of resources by the local community was likely lower due to the decline in zooplankton biomass. In contrast, phytoplankton arrived at a community where the biomass of their competitors was reduced, and the heatwave even contributed to a successful establishment by suppressing their consumers and thereby releasing available nutrients.

Global climate warming is accelerating with more frequent and severe heatwaves in both aquatic (Woolway et al., 2021) and terrestrial realms (Perkins-Kirkpatrick & Lewis, 2020), with negative consequences for biodiversity, ecosystem functioning, and services. Our results illustrate that a relatively short, ca. 10-day-long heatwave can already alter the main pathways of energy flow in aquatic ecosystems through differential responses across trophic groups. We also showed that dispersal from a regional species pool can enhance community resilience by faster biomass recovery at the level of primary producers, which may contribute to more stable levels of total community biomass in the long term. Our findings have important implications for conservation planning under future climate change. As a consequence of accelerating habitat loss and fragmentation, aquatic habitats become spatially more isolated (Davidson & Davidson, 2014; Hassall, 2014; Horváth et al., 2019), and consequently less buffered against heatwave effects under the future climates. Conservation actions therefore need to focus on protecting habitat networks and maintaining sufficiently diverse metacommunities, as connectivity among local habitats is likely to be essential for their resilience.

## Acknowledgements

This work was supported through the Transnational Access Programme of the AQUACOSM project, which has received funding from the European Union’s Horizon 2020 research and innovation programme under grant agreement No. 731065 (Transnational Access projects: MicroDISCO, CoDRes, AQUADOMES, Meso_Zoo, ZooDispersAlp). CFV acknowledges further support by the AQUACOSM-plus project of the European Union’s Horizon 2020 research and innovation programme under grant agreement No. 871081 and by the RRF-2.3.1-21-2022-00014 project. The authors thank the firemen of Lunz am See for their help in filling the mesocosms with lake water, as well as Marina Ivanković, Robert Fischer, Zsuzsanna Márton, Iris Schachner, Beate Pitzl, Claudia Schneider, and Zsófia Horváth for their practical help during the experimental and laboratory works.

## Author contributions

CFV and RP conceived the main study idea. All authors contributed to practical works during the experiment, including sampling, sample processing, and laboratory works. AA performed the analysis (estimation of abundance, taxonomic identification) of phytoplankton samples, while LS performed the analysis of zooplankton with the help of CFV. AH-E and CFV performed the statistical analyses. All authors contributed to scientific discussions during the Transnational Access. CFV wrote the first draft of the manuscript after which all authors contributed with feedback and edits.

## References

Adrian, R., & Schneider-Olt, B. (1999). Top-down effects of crustacean zooplankton on pelagic microorganisms in a mesotrophic lake. Journal of Plankton Research, 21(11), 2175–2190. https://doi.org/10.1093/plankt/21.11.2175

Anderson, M. J. (2001). A new method for non-parametric multivariate analysis of variance. Austral Ecology, 26(1), 32–46. https://doi.org/10.1111/j.1442-9993.2001.01070.pp.x

Arar, E., & Collins, G. (1997). Method 445.0. In-vitro determination of chlorophyll a and pheophytin a in marine and freshwater algae by fluorescence. US Environmental Protection Agency, National Exposure Research Laboratory, Office of Research and Development. https://cfpub.epa.gov/si/si_public_record_report.cfm?Lab=NERL&dirEntryId=309417

Bartoń, K. (2022). MuMIn: Multi-Model Inference (1.46.0). https://CRAN.R-project.org/package=MuMIn

Bender, E. A., Case, T. J., & Gilpin, M. E. (1984). Perturbation experiments in community ecology: Theory and practice. Ecology, 65(1), 1–13. https://doi.org/10.2307/1939452

Bennett, J. M., Sunday, J., Calosi, P., Villalobos, F., Martínez, B., Molina-Venegas, R., Araújo, M. B., Algar, A. C., Clusella-Trullas, S., Hawkins, B. A., Keith, S. A., Kühn, I., Rahbek, C., Rodríguez, L., Singer, A., Morales-Castilla, I., & Olalla-Tárraga, M. Á. (2021). The evolution of critical thermal limits of life on Earth. Nature Communications, 12(1), 1198. https://doi.org/10.1038/s41467-021-21263-8

Bottrell, H. H., Duncan, A., Gliwicz, Z., Grygierek, E., Herzig, A., Hilbricht-Ilkowska, A., Kurasawa, H., Larsson, P., & Weglenska, T. (1976). Review of some problems in zooplankton production studies. Norwegian Journal of Zoology, 21, 477–483.

Brandl, Z. (2005). Freshwater copepods and rotifers: Predators and their prey. Hydrobiologia, 546(1), 475–489. https://doi.org/10.1007/s10750-005-4290-3

Brett, M. T., Müller-Navarra, D. C., & Persson, J. (2009). Crustacean zooplankton fatty acid composition. In M. Kainz, M. T. Brett, & M. T. Arts (Eds.), Lipids in Aquatic Ecosystems (pp. 115–146). Springer. https://doi.org/10.1007/978-0-387-89366-2_6

Brown, J. H., Gillooly, J. F., Allen, A. P., Savage, V. M., & West, G. B. (2004). Toward a metabolic theory of ecology. Ecology, 85(7), 1771–1789. https://doi.org/10.1890/03-9000

Clark, G. F., & Johnston, E. L. (2011). Temporal change in the diversity–invasibility relationship in the presence of a disturbance regime. Ecology Letters, 14(1), 52–57. https://doi.org/10.1111/j.1461-0248.2010.01550.x

Clesceri, L. S., Greenberg, A. E., & Eaton, A. D. (Eds.). (1999). Standard Methods for the Examination of Water and Wastewater (20th ed). APHA, AWWA, WEF.

Comte, L., & Olden, J. D. (2017). Climatic vulnerability of the world’s freshwater and marine fishes. Nature Climate Change, 7(10), 718–722. https://doi.org/10.1038/nclimate3382

Davidson, K., Roberts, E. C., & Gilpin, L. C. (2002). The relationship between carbon and biovolume in marine microbial mesocosms under different nutrient regimes. European Journal of Phycology, 37(4), 501–507. https://doi.org/10.1017/S096702620200389X

Davidson, N. C., & Davidson, N. C. (2014). How much wetland has the world lost? Long-term and recent trends in global wetland area. Marine and Freshwater Research, 65(10), 934–941. https://doi.org/10.1071/MF14173

Davis, M. A., Grime, J. P., & Thompson, K. (2000). Fluctuating resources in plant communities: A general theory of invasibility. Journal of Ecology, 88(3), 528–534. https://doi.org/10.1046/j.1365-2745.2000.00473.x

Davison, R., Jacquemyn, H., Adriaens, D., Honnay, O., Kroon, H. D., & Tuljapurkar, S. (2010). Demographic effects of extreme weather events on a short-lived calcareous grassland species: Stochastic life table response experiments. Journal of Ecology, 98(2), 255–267. https://doi.org/10.1111/j.1365-2745.2009.01611.x

de Boer, M. K., Moor, H., Matthiessen, B., Hillebrand, H., & Eriksson, B. K. (2014). Dispersal restricts local biomass but promotes the recovery of metacommunities after temperature stress. Oikos, 123(6), 762–768. https://doi.org/10.1111/j.1600-0706.2013.00927.x

Dlouhá, Š., Thielsch, A., Kraus, R. H. S., Seda, J., Schwenk, K., & Petrusek, A. (2010). Identifying hybridizing taxa within the Daphnia longispina species complex: A comparison of genetic methods and phenotypic approaches. Hydrobiologia, 643(1), 107–122. https://doi.org/10.1007/s10750-010-0128-8

Duffy, J. E., Cardinale, B. J., France, K. E., McIntyre, P. B., Thébault, E., & Loreau, M. (2007). The functional role of biodiversity in ecosystems: Incorporating trophic complexity. Ecology Letters, 10(6), 522–538. https://doi.org/10.1111/j.1461-0248.2007.01037.x

Eggers, S. L., Eriksson, B. K., & Matthiessen, B. (2012). A heat wave and dispersal cause dominance shift and decrease biomass in experimental metacommunities. Oikos, 121(5), 721–733. https://doi.org/10.1111/j.1600-0706.2011.19714.x

Fischer, E. M., & Schär, C. (2010). Consistent geographical patterns of changes in high-impact European heatwaves. Nature Geoscience, 3(6), 398–403. https://doi.org/10.1038/ngeo866

Frölicher, T. L., Fischer, E. M., & Gruber, N. (2018). Marine heatwaves under global warming. Nature, 560(7718), 360–364. https://doi.org/10.1038/s41586-018-0383-9

Fussmann, K. E., Schwarzmüller, F., Brose, U., Jousset, A., & Rall, B. C. (2014). Ecological stability in response to warming. Nature Climate Change, 4(3), 206–210. https://doi.org/10.1038/nclimate2134

Gerhard, M., Schlenker, A., Hillebrand, H., & Striebel, M. (2022). Environmental stoichiometry mediates phytoplankton diversity effects on communities’ resource use efficiency and biomass. Journal of Ecology, 110(2), 430–442. https://doi.org/10.1111/1365-2745.13811

Guelzow, N., Muijsers, F., Ptacnik, R., & Hillebrand, H. (2017). Functional and structural stability are linked in phytoplankton metacommunities of different connectivity. Ecography, 40(6), 719–732. https://doi.org/10.1111/ecog.02458

Gutschick, V. P., & BassiriRad, H. (2003). Extreme events as shaping physiology, ecology, and evolution of plants: Toward a unified definition and evaluation of their consequences. New Phytologist, 160(1), 21–42. https://doi.org/10.1046/j.1469-8137.2003.00866.x

Haegeman, B., & Loreau, M. (2014). General relationships between consumer dispersal, resource dispersal and metacommunity diversity. Ecology Letters, 17(2), 175–184. https://doi.org/10.1111/ele.12214

Hansen, H. P., & Koroleff, F. (1999). Determination of nutrients. In K. Grasshoff, K. Kremling, & M. Ehrhardt (Eds.), Methods of Seawater Analysis (3rd ed., pp. 159–228). Wiley-VCH. https://doi.org/10.1002/9783527613984.ch10

Harris, R. M. B., Beaumont, L. J., Vance, T. R., Tozer, C. R., Remenyi, T. A., Perkins-Kirkpatrick, S. E., Mitchell, P. J., Nicotra, A. B., McGregor, S., Andrew, N. R., Letnic, M., Kearney, M. R., Wernberg, T., Hutley, L. B., Chambers, L. E., Fletcher, M.-S., Keatley, M. R., Woodward, C. A., Williamson, G., …Bowman, D. M. J. S. (2018). Biological responses to the press and pulse of climate trends and extreme events. Nature Climate Change, 8(7), 579–587. https://doi.org/10.1038/s41558-018-0187-9

Hassall, C. (2014). The ecology and biodiversity of urban ponds. Wiley Interdisciplinary Reviews: Water, 1(2), 187–206. https://doi.org/10.1002/wat2.1014

Hillebrand, H., Dürselen, C.-D., Kirschtel, D., Pollingher, U., & Zohary, T. (1999). Biovolume calculation for pelagic and benthic microalgae. Journal of Phycology, 35(2), 403–424. https://doi.org/10.1046/j.1529-8817.1999.3520403.x

Hodgson, D., McDonald, J. L., & Hosken, D. J. (2015). What do you mean, ‘resilient’? Trends in Ecology & Evolution, 30(9), 503–506. https://doi.org/10.1016/j.tree.2015.06.010

Horváth, Z., Ptacnik, R., Vad, C. F., & Chase, J. M. (2019). Habitat loss over six decades accelerates regional and local biodiversity loss via changing landscape connectance. Ecology Letters, 22(6), 1019–1027. https://doi.org/10.1111/ele.13260

Huber, V., Adrian, R., & Gerten, D. (2010). A matter of timing: Heat wave impact on crustacean zooplankton. Freshwater Biology, 55(8), 1769–1779. https://doi.org/10.1111/j.1365-2427.2010.02411.x

Ingrisch, J., & Bahn, M. (2018). Towards a comparable quantification of resilience. Trends in Ecology & Evolution, 33(4), 251–259. https://doi.org/10.1016/j.tree.2018.01.013

IPCC. (2021). Summary for Policymakers. In V. Masson-Delmotte, P. Zhai, A. Pirani, S. L. Connors, C. Péan, S. Berger, N. Caud, Y. Chen, L. Goldfarb, M. I. Gomis, M. Huang, K. Leitzell, E. Lonnoy, J. B. R. Matthews, T. K. Maycock, T. Waterfield, O. Yelekçi, R. Yu, & B. Zhou (Eds.), Climate Change 2021: The Physical Science Basis. Contribution of Working Group I to the Sixth Assessment Report of the Intergovernmental Panel on Climate Change (pp. 3–32). Cambridge University Press.

Jentsch, A., Kreyling, J., & Beierkuhnlein, C. (2007). A new generation of climate-change experiments: Events, not trends. Frontiers in Ecology and the Environment, 5(7), 365– 374. https://doi.org/10.1890/1540-9295(2007)5[365:ANGOCE]2.0.CO;2

Jiguet, F., Julliard, R., Thomas, C. D., Dehorter, O., Newson, S. E., & Couvet, D. (2006). Thermal range predicts bird population resilience to extreme high temperatures. Ecology Letters, 9(12), 1321–1330. https://doi.org/10.1111/j.1461-0248.2006.00986.x

Koste, W. (1978). Rotatoria. Die Rädertiere Mitteleuropas. 1 Textband (2nd ed.). Gebrüder Bornträger.

Kratina, P., Rosenbaum, B., Gallo, B., Horas, E. L., & O’Gorman, E. J. (2022). The combined effects of warming and body size on the stability of predator-prey interactions. Frontiers in Ecology and Evolution, 9, 772078.

Lampert, W., Fleckner, W., Rai, H., & Taylor, B. E. (1986). Phytoplankton control by grazing zooplankton: A study on the spring clear-water phase. Limnology and Oceanography, 31(3), 478–490. https://doi.org/10.4319/lo.1986.31.3.0478

Lampert, W., & Muck, P. (1985). Multiple aspects of food limitation in zooplankton communities: The Daphnia - Eudiaptomus example. Ergebnisse Der Limnologie/Advances in Limnology, 21, 311–322.

Leibold, M. A., Chase, J. M., & Ernest, S. K. M. (2017). Community assembly and the functioning of ecosystems: How metacommunity processes alter ecosystems attributes. Ecology, 98(4), 909–919. https://doi.org/10.1002/ecy.1697

Limberger, R., Pitt, A., Hahn, M. W., & Wickham, S. A. (2019). Spatial insurance in multitrophic metacommunities. Ecology Letters, 22(11), 1828–1837. https://doi.org/10.1111/ele.13365

López-Urrutia, Á., San Martin, E., Harris, R. P., & Irigoien, X. (2006). Scaling the metabolic balance of the oceans. Proceedings of the National Academy of Sciences, 103(23), 8739–8744. https://doi.org/10.1073/pnas.0601137103

Loreau, M., Mouquet, N., & Gonzalez, A. (2003). Biodiversity as spatial insurance in heterogeneous landscapes. Proceedings of the National Academy of Sciences, 100(22), 12765–12770. https://doi.org/10.1073/pnas.2235465100

Ludovisi, A., Todini, C., Pandolfi, P., & Taticchi, M. I. (2008). Scale patterns of diel distribution of the copepod Cyclops abyssorum Sars in a regulated lake: The relative importance of physical and biological factors. Journal of Plankton Research, 30(5), 495–509. https://doi.org/10.1093/plankt/fbn017

Lund, J. W. G., Kipling, C., & Le Cren, E. D. (1958). The inverted microscope method of estimating algal numbers and the statistical basis of estimations by counting. Hydrobiologia, 11(2), 143–170. https://doi.org/10.1007/BF00007865

Matthiessen, B., & Hillebrand, H. (2006). Dispersal frequency affects local biomass production by controlling local diversity. Ecology Letters, 9(6), 652–662. https://doi.org/10.1111/j.1461-0248.2006.00916.x

McCauley, E. (1984). The estimation of the abundance and biomass of zooplankton in samples. In J. A. Downing & F. H. Rigler (Eds.), A manual on methods for the assessment of secondary productivity in fresh waters. (2nd ed., Vol. 17, pp. 228–265). Blackwell.

McGlinn, D. J., Xiao, X., May, F., Gotelli, N. J., Engel, T., Blowes, S. A., Knight, T. M., Purschke, O., Chase, J. M., & McGill, B. J. (2019). Measurement of Biodiversity (MoB): A method to separate the scale-dependent effects of species abundance distribution, density, and aggregation on diversity change. Methods in Ecology and Evolution, 10(2), 258– 269. https://doi.org/10.1111/2041-210X.13102

McGlinn, D., Xiao, X., McGill, B., May, F., Engel, T., Oliver, C., Blowes, S., Knight, T., Purschke, O., Gotelli, N., & Chase, J. (2021). mobr: Measurement of Biodiversity (2.0.2). https://CRAN.R-project.org/package=mobr

O’Connor, M. I., Gilbert, B., & Brown, C. J. (2011). Theoretical predictions for how temperature affects the dynamics of interacting herbivores and plants. The American Naturalist, 178(5), 626–638. https://doi.org/10.1086/662171

O’Connor, M. I., Piehler, M. F., Leech, D. M., Anton, A., & Bruno, J. F. (2009). Warming and resource availability shift food web structure and metabolism. PLOS Biology, 7(8), e1000178. https://doi.org/10.1371/journal.pbio.1000178

Oksanen, J., Blanchet, F. G., Friendly, M., Kindt, R., Legendre, P., McGlinn, D., Minchin, P. R., O’Hara, R. B., Simpson, G. L., Solymos, P., Stevens, M. H. H., Szoecs, E., & Wagner, H. (2020). vegan: Community Ecology Package (2.5-7). https://CRAN.R-project.org/package=vegan

Oliver, E. C. J., Donat, M. G., Burrows, M. T., Moore, P. J., Smale, D. A., Alexander, L. V., Benthuysen, J. A., Feng, M., Sen Gupta, A., Hobday, A. J., Holbrook, N. J., Perkins-Kirkpatrick, S. E., Scannell, H. A., Straub, S. C., & Wernberg, T. (2018). Longer and more frequent marine heatwaves over the past century. Nature Communications, 9(1), 1324. https://doi.org/10.1038/s41467-018-03732-9

Perkins-Kirkpatrick, S. E., & Lewis, S. C. (2020). Increasing trends in regional heatwaves. Nature Communications, 11(1), 3357. https://doi.org/10.1038/s41467-020-16970-7

Persson, J., Brett, M. T., Vrede, T., & Ravet, J. L. (2007). Food quantity and quality regulation of trophic transfer between primary producers and a keystone grazer (Daphnia) in pelagic freshwater food webs. Oikos, 116(7), 1152–1163. https://doi.org/10.1111/j.0030-1299.2007.15639.x

Petchey, O. L., McPhearson, P. T., Casey, T. M., & Morin, P. J. (1999). Environmental warming alters food-web structure and ecosystem function. Nature, 402(6757), 69–72. https://doi.org/10.1038/47023

Pinheiro, J. C., & Bates, D. M. (2000). Mixed-Effects Models in S and S-PLUS (1st ed.). Springer-Verlag. https://link.springer.com/book/10.1007/b98882

Pinsky, M. L., Eikeset, A. M., McCauley, D. J., Payne, J. L., & Sunday, J. M. (2019). Greater vulnerability to warming of marine versus terrestrial ectotherms. Nature, 569(7754), 108–111. https://doi.org/10.1038/s41586-019-1132-4

Poxleitner, M., Trommer, G., Lorenz, P., & Stibor, H. (2016). The effect of increased nitrogen load on phytoplankton in a phosphorus-limited lake. Freshwater Biology, 61(11), 1966–1980. https://doi.org/10.1111/fwb.12829

Purvis, A., Gittleman, J. L., Cowlishaw, G., & Mace, G. M. (2000). Predicting extinction risk in declining species. Proceedings of the Royal Society of London. Series B: Biological Sciences, 267(1456), 1947–1952. https://doi.org/10.1098/rspb.2000.1234

R Core Team. (2020). R: A Language and Environment for Statistical Computing. (4.0.2). R Foundation for Statistical Computing. URL https://www.R-project.org/

Rall, B. C., Vucic-Pestic, O., Ehnes, R. B., Emmerson, M., & Brose, U. (2010). Temperature, predator–prey interaction strength and population stability. Global Change Biology, 16(8), 2145–2157. https://doi.org/10.1111/j.1365-2486.2009.02124.x

Reiss, J., & Schmid-Araya, J. M. (2008). Existing in plenty: Abundance, biomass and diversity of ciliates and meiofauna in small streams. Freshwater Biology, 53(4), 652–668. https://doi.org/10.1111/j.1365-2427.2007.01907.x

Romero, G. Q., Gonçalves-Souza, T., Kratina, P., Marino, N. A. C., Petry, W. K., Sobral-Souza, T., & Roslin, T. (2018). Global predation pressure redistribution under future climate change. Nature Climate Change, 8(12), 1087–1091. https://doi.org/10.1038/s41558-018-0347-y

Ross, S. R. P.-J., García Molinos, J., Okuda, A., Johnstone, J., Atsumi, K., Futamura, R., Williams, M. A., Matsuoka, Y., Uchida, J., Kumikawa, S., Sugiyama, H., Kishida, O., & Donohue, I. (2022). Predators mitigate the destabilising effects of heatwaves on multitrophic stream communities. Global Change Biology, 28(2), 403–416. https://doi.org/10.1111/gcb.15956

Sentis, A., Hemptinne, J.-L., & Brodeur, J. (2013). Effects of simulated heat waves on an experimental plant–herbivore–predator food chain. Global Change Biology, 19(3), 833–842. https://doi.org/10.1111/gcb.12094

Shurin, J. B., Clasen, J. L., Greig, H. S., Kratina, P., & Thompson, P. L. (2012). Warming shifts top-down and bottom-up control of pond food web structure and function. Philosophical Transactions of the Royal Society B: Biological Sciences, 367(1605), 3008–3017. https://doi.org/10.1098/rstb.2012.0243

Slobodkin, L. B. (1959). Energetics in Daphnia pulex populations. Ecology, 40(2), 232–243. https://doi.org/10.2307/1930033

Smale, D. A., & Wernberg, T. (2013). Extreme climatic event drives range contraction of a habitat-forming species. Proceedings of the Royal Society B: Biological Sciences, 280(1754), 20122829. https://doi.org/10.1098/rspb.2012.2829

Smale, D. A., Wernberg, T., Oliver, E. C. J., Thomsen, M., Harvey, B. P., Straub, S. C., Burrows, M. T., Alexander, L. V., Benthuysen, J. A., Donat, M. G., Feng, M., Hobday, A. J., Holbrook, N. J., Perkins-Kirkpatrick, S. E., Scannell, H. A., Sen Gupta, A., Payne, B. L., & Moore, P. J. (2019). Marine heatwaves threaten global biodiversity and the provision of ecosystem services. Nature Climate Change, 9(4), 306–312. https://doi.org/10.1038/s41558-019-0412-1

Stillman, J. H. (2019). Heat waves, the new normal: Summertime temperature extremes will impact animals, ecosystems, and human communities. Physiology, 34(2), 86–100. https://doi.org/10.1152/physiol.00040.2018

Striebel, M., Kirchmaier, L., & Hingsamer, P. (2013). Different mixing techniques in experimental mesocosms—Does mixing affect plankton biomass and community composition? Limnology and Oceanography: Methods, 11(4), 176–186. https://doi.org/10.4319/lom.2013.11.176

Sunday, J. M., Bates, A. E., & Dulvy, N. K. (2012). Thermal tolerance and the global redistribution of animals. Nature Climate Change, 2(9), 686–690. https://doi.org/10.1038/nclimate1539

Symons, C. C., & Arnott, S. E. (2013). Regional zooplankton dispersal provides spatial insurance for ecosystem function. Global Change Biology, 19(5), 1610–1619. https://doi.org/10.1111/gcb.12122

Symons, C. C., & Arnott, S. E. (2014). Timing is everything: Priority effects alter community invasibility after disturbance. Ecology and Evolution, 4(4), 397–407. https://doi.org/10.1002/ece3.940

Thébault, E., & Loreau, M. (2003). Food-web constraints on biodiversity–ecosystem functioning relationships. Proceedings of the National Academy of Sciences, 100(25), 14949–14954. https://doi.org/10.1073/pnas.2434847100

Thompson, P. L., Beisner, B. E., & Gonzalez, A. (2015). Warming induces synchrony and destabilizes experimental pond zooplankton metacommunities. Oikos, 124(9), 1171– 1180. https://doi.org/10.1111/oik.01945

Thompson, P. L., & Gonzalez, A. (2017). Dispersal governs the reorganization of ecological networks under environmental change. Nature Ecology & Evolution, 1(6), 1–8. https://doi.org/10.1038/s41559-017-0162

Thompson, P. L., Rayfield, B., & Gonzalez, A. (2017). Loss of habitat and connectivity erodes species diversity, ecosystem functioning, and stability in metacommunity networks. Ecography, 40(1), 98–108. https://doi.org/10.1111/ecog.02558

Thompson, P. L., & Shurin, J. B. (2012). Regional zooplankton biodiversity provides limited buffering of pond ecosystems against climate change. Journal of Animal Ecology, 81(1), 251–259. https://doi.org/10.1111/j.1365-2656.2011.01908.x

Thompson, R. M., Beardall, J., Beringer, J., Grace, M., & Sardina, P. (2013). Means and extremes: Building variability into community-level climate change experiments. Ecology Letters, 16(6), 799–806. https://doi.org/10.1111/ele.12095

Utermöhl, H. (1958). Zur vervollkommnung der quantitativen phytoplankton-methodik. Mitteilungen Internationale Vereinigung für theoretische und angewandte Limnologie, 9, 1–38.

Vadstein, O., Jensen, A., Olsen, Y., & Reinertsen, H. (1988). Growth and phosphorus status of limnetic phytoplankton and bacteria. Limnology and Oceanography, 33, 489–503.

Vasseur, D. A., DeLong, J. P., Gilbert, B., Greig, H. S., Harley, C. D. G., McCann, K. S., Savage, V., Tunney, T. D., & O’Connor, M. I. (2014). Increased temperature variation poses a greater risk to species than climate warming. Proceedings of the Royal Society B: Biological Sciences, 281(1779), 20132612. https://doi.org/10.1098/rspb.2013.2612

Velthuis, M., de Senerpont Domis, L. N., Frenken, T., Stephan, S., Kazanjian, G., Aben, R., Hilt, S., Kosten, S., van Donk, E., & Van de Waal, D. B. (2017). Warming advances top-down control and reduces producer biomass in a freshwater plankton community. Ecosphere, 8(1), e01651. https://doi.org/10.1002/ecs2.1651

Wernberg, T., Smale, D. A., Tuya, F., Thomsen, M. S., Langlois, T. J., de Bettignies, T., Bennett, S., & Rousseaux, C. S. (2013). An extreme climatic event alters marine ecosystem structure in a global biodiversity hotspot. Nature Climate Change, 3(1), 78–82. https://doi.org/10.1038/nclimate1627

Wickham, H., Chang, W., Henry, L., Pedersen, T. L., Takahashi, K., Wilke, C., Woo, K., Yutani, H., Dunnington, D., & RStudio. (2021). ggplot2: Create Elegant Data Visualisations Using the Grammar of Graphics (3.3.5). https://CRAN.R-project.org/package=ggplot2

Woodward, G., Bonada, N., Brown, L. E., Death, R. G., Durance, I., Gray, C., Hladyz, S., Ledger, M. E., Milner, A. M., Ormerod, S. J., Thompson, R. M., & Pawar, S. (2016). The effects of climatic fluctuations and extreme events on running water ecosystems. Philosophical Transactions ofV the Royal Society B: Biological Sciences, 371(1694), 20150274. https://doi.org/10.1098/rstb.2015.0274

Woolway, R. I., Jennings, E., Shatwell, T., Golub, M., Pierson, D. C., & Maberly, S. C. (2021). Lake heatwaves under climate change. Nature, 589(7842), 402–407. https://doi.org/10.1038/s41586-020-03119-1

Yampolsky, L. Y., Zeng, E., Lopez, J., Williams, P. J., Dick, K. B., Colbourne, J. K., & Pfrender, M. E. (2014). Functional genomics of acclimation and adaptation in response to thermal stress in Daphnia. BMC Genomics, 15(1), 859. https://doi.org/10.1186/1471-2164-15-859

Zhang, H., Urrutia-Cordero, P., He, L., Geng, H., Chaguaceda, F., Xu, J., & Hansson, L.-A. (2018). Life-history traits buffer against heat wave effects on predator–prey dynamics in zooplankton. Global Change Biology, 24(10), 4747–4757. https://doi.org/10.1111/gcb.14371

Zhang, P., Leeuwen, C. H. A. van, Bogers, D., Poelman, M., Xu, J., & Bakker, E. S. (2020). Ectothermic omnivores increase herbivory in response to rising temperature. Oikos, 129(7), 1028–1039. https://doi.org/10.1111/oik.07082

