## Supplementary material for "Dispersal provides trophic-level dependent insurance against a heatwave in freshwater ecosystems"

### **Table S1 –** List of lakes sampled to obtain the dispersal inoculum for the experiment.

| **Lake** | **Coordinates** | **Max. depth (m)** |
| --- | --- | --- |
| Grabensee | N 47°59'28" E 13°05'46" | 14 |
| Obertrummersee | N 47°57'40" E 13°04'50" | 36 |
| Mattsee | N 47°59'10" E 13°07'30" | 42 |
| Irrsee | N 47°54'44" E 13°18'25" | 32 |
| Mondsee | N 47°49'36" E 13°22'48" | 68 |
| Obinger See | N 48°00'11" E 12°24'56" | 45 |
| Brunnsee | N 47°59'03" E 12°26'11" | 20 |
| Griessee | N 47°59'11" E 12°26'33" | 12 |
| Seeleitensee | N 47°58'32" E 12°25'56" | 15 |
| Klostersee | N 47°58'25" E 12°27'03" | 16 |
| Bansee | N 47°57'51" E 12°26'25" | 4 |
| Eschenauer See | N 47°56'41" E 12°23'51" | 3 |
| Pelhamer See | N 47°56'01" E 12°20'54" | 21 |
| Hartsee | N 47°55'35" E 12°21'54" | 39 |
| Langbürgenersee | N 47°54'08" E 12°20'51" | 37 |
| Thalersee | N 47°54'16" E 12°20'17" | 47 |
| Chiemsee | N 47°53'50" E 12°27'49" | 73 |
| Simsee | N 47°52'03" E 12°13'48" | 23 |

### **Table S2** – List of zooplankton taxa identified in the study. Species that were absent in the local (i.e., Lake Lunz) community, and were only introduced with the regional dispersal inoculum are marked as ‘regional taxa’. Taxa that were present in both the local and regional species pool are marked with an asterisk. Besides, the occurrence of each species at the focal time points is indicated (present: ‘+’, absent: ‘-‘). We note that identification was based on phenotype, and therefore, we were not able to distinguish between different genotypes of morphologically cryptic lineages, which may have been introduced by dispersal.

| **Taxa** | **Regional taxa** | **Presence at t_1_** | **Presence at t_2_** | **Presence at t_4_** |
| --- | --- | --- | --- | --- |
| *Alona rectangula* |  | **-** | **+** | **+** |
| *Asplanchna* cf. *priodonta* |  | **+** | **-** | **+** |
| *Bosmina longispina** |  | **+** | **+** | **+** |
| *Asplanchna* cf. herricki | **x** | **-** | **-** | **-** |
| *Ceriodaphnia* sp. | **x** | **-** | **-** | **-** |
| *Chydorus sphaericus* |  | **+** | **+** | **+** |
| *Cyclops abyssorum* |  | **+** | **+** | **+** |
| *Daphnia cucullata* | **x** | **+** | **-** | **+** |
| *Daphnia* cf. *longispina** |  | **+** | **+** | **+** |
| *Diaphanosoma brachyurum* | **x** | **-** | **-** | **-** |
| *Euchlanis* cf. *dilatata* |  | **+** | **+** | **+** |
| *Eudiaptomus gracilis** |  | **+** | **+** | **+** |
| *Kellicottia longispina* |  | **+** | **-** | **-** |
| *Keratella cochlearis* |  | **+** | **+** | **+** |
| *Keratella quadrata* |  | **+** | **-** | **-** |
| *Mesocyclops leuckarti* | **x** | **+** | **+** | **+** |
| *Microcyclops rubellus* |  | **-** | **-** | **+** |
| *Polyarthra vulgaris* |  | **+** | **+** | **-** |
| *Synchaeta* cf. *longipes* |  | **+** | **+** | **+** |
| *Thermocyclops crassus* | **x** | **+** | **+** | **+** |
| *Thermocyclops oithonoides* | **x** | **+** | **+** | **-** |

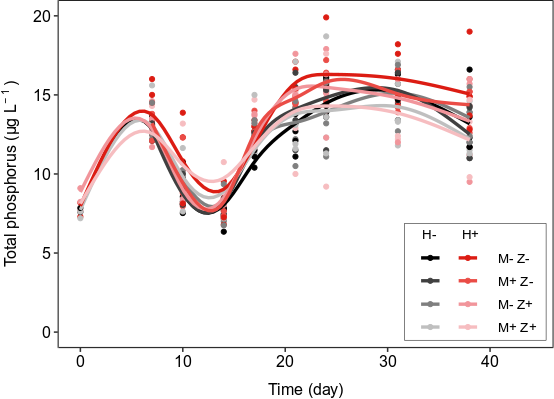

### **Figure S1** – Concentrations of total phosphorus over the experimental duration. Solid lines represent smoothed conditional mean curves based on generalised additive models (GAM) to visualise dynamics in the different treatments (heatwave: H-, H+, dispersal of microorganisms: M-, M+, dispersal of zooplankton: Z-, Z+). N = 5 for all treatment combinations per day. GAM models were fit with the *stat_smooth* function (using mgcv gam fitting and formula: y~s(x,k=8)) of the ggplot2 R package.

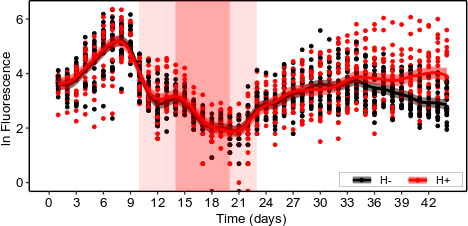

### **Figure S2** – Pattern of daily chlorophyll *a* in vivo fluorescence over the experimental duration coloured according to the heatwave (H+) and control (H-) treatments. Red-coloured background shading denotes the time interval of the simulated heatwave in H+ treatments (light shading: heating and cooling phases, dark shading: culmination phase with a +5^o^C offset in H+ vs H-). Solid lines represent fitted generalised additive models (with 95% confidence intervals) illustrating dynamics in the H- and H+ treatments (N = 20 per treatment and per day).

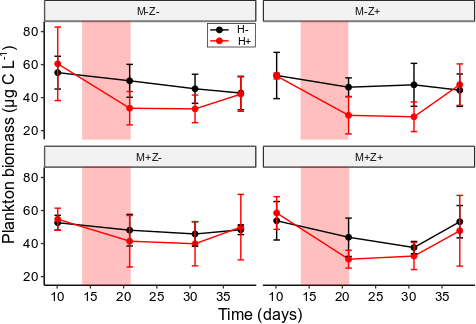

### **Figure S3** – Total plankton biomass (expressed in carbon mass, mean ± SD), i.e., the sum of zooplankton carbon biomass and POC at the four sampling days. N = 5 for all treatment combinations per day. Red shading denotes the time interval of the culmination phase (+5^o^C offset in H+ vs H-) of the heatwave in H+. Results of the linear models testing for resistance (i.e., biomass difference between t_1_ and t_2_) and resilience (i.e., biomass difference between t_3_ and t_4_) are presented in Table 1.

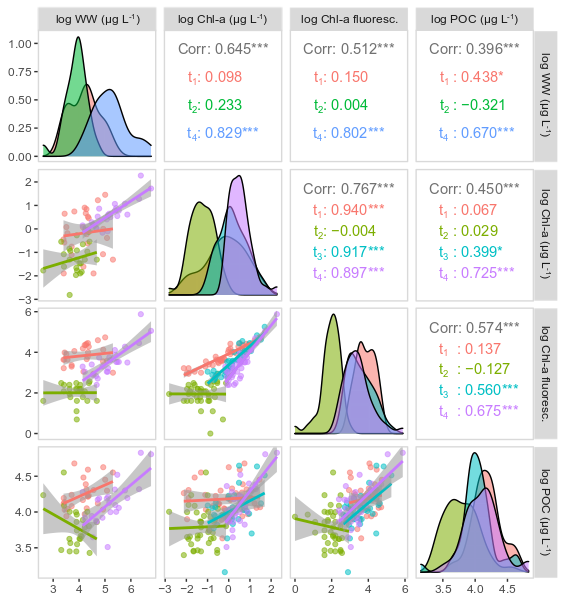

### **Figure S4** – Pairwise correlation plot of POC, phytoplankton wet weight (WW) and Chl-a (both concentration and fluorescence). The relationships were tested by Pearson’s correlations among variables. All data were log-transformed prior to analyses. The correlation coefficients are presented for the pooled dataset (i.e., data of all sampling dates analysed together) and separately for individual focal sampling dates (indicated by different colours). Histograms illustrate the distribution of data. Significance levels: ‘*’ p<0.05, ‘**’ p<0.01, ‘***’ p<0.001.

### **Table S3** – Summary statistics (estimated parameters, standard error and p values) of multiple linear regression models testing resistance and resilience and their interaction with dispersal (i.e., zooplankton and microorganisms) in Cladocera and Copepoda, the two dominant groups of zooplankton in this study. Resistance is defined as the biomass change (expressed in carbon) during the heatwave (i.e., difference between t_1_ and t_2_), while resilience as the change in the after-heatwave period (i.e., difference between t_3_ and t_4_). Abbreviations: ClaB – Cladocera biomass, CopB – Copepoda biomass, initial – mean-centred initial biomass. Treatment labels: H+: heatwave, M+: dispersal of microorganisms, Z-: dispersal of zooplankton. Significant results (p<0.05) are highlighted with bold letters, while marginally significant ones (p<0.1) with italics.

|  | **ClaB** | | |  | **CopB** | | |
| --- | --- | --- | --- | --- | --- | --- | --- |
|  | est. | SE | *p* |  | est. | SE | *p* |
| **Resistance** | | | | | | | |
| intercept | **12.1** | **3.38** | **0.001** |  | **3.7** | **1.15** | **0.038** |
| initial | **-1.3** | **0.15** | **<0.001** |  | **-0.6** | **0.16** | **<0.001** |
| H+ | **-21.3** | **4.79** | **<0.001** |  | *-3.1* | *1.64* | *0.073* |
| M+Z- | -6.9 | 4.88 | 0.166 |  | -1.4 | 1.65 | 0.390 |
| M-Z+ | *-9.2* | 4.86 | *0.068* |  | -2.4 | 1.64 | 0.158 |
| M+Z+ | **-12.3** | **4.78** | **0.015** |  | -1.4 | 1.68 | 0.309 |
| H+ × M+Z- | 3.5 | 6.82 | 0.609 |  | 2.3 | 2.36 | 0.335 |
| H+ × M-Z+ | 4.3 | 6.76 | 0.533 |  | 1.3 | 2.33 | 0.576 |
| H+ × M+Z+ | 7.8 | 6.81 | 0.261 |  | 1.7 | 2.34 | 0.471 |
| **R^2^** | 0.81 | | |  | 0.34 | | |
| **Recovery** | | | | | | | |
| intercept | 4.52 | 4.20 | 0.290 |  | 1.18 | 0.72 | 0.115 |
| initial | *0.34* | 0.18 | *0.076* |  | 0.05 | 0.10 | 0.621 |
| H+ | 0.85 | 5.94 | 0.887 |  | 0.95 | 1.02 | 0.359 |
| M+Z- | -2.91 | 6.05 | 0.634 |  | 0.49 | 1.03 | 0.640 |
| M-Z+ | 9.30 | 6.03 | 0.133 |  | 1.40 | 1.03 | 0.184 |
| M+Z+ | 1.52 | 5.93 | 0.800 |  | 0.53 | 1.05 | 0.620 |
| H+ × M+Z- | 4.02 | 8.45 | 0.638 |  | -1.49 | 1.48 | 0.320 |
| H+ × M-Z+ | -9.63 | 8.38 | 0.259 |  | -0.58 | 1.46 | 0.691 |
| H+ × M+Z+ | -0.60 | 8.45 | 0.944 |  | -1.20 | 1.46 | 0.420 |
| **R^2^** | 0.00 | | |  | 0.00 | | |

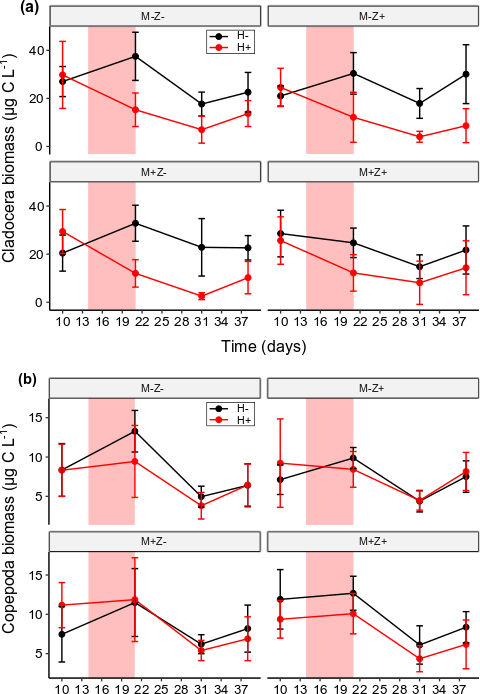

**Figure S5** – Zooplankton biomass (carbon mass, mean ± SD) at the four sampling days shown separately for **(a)** Cladocera and **(b)** Copepoda. N = 5 for all treatment combinations per day. Red shading denotes the time interval of the culmination phase (+5^o^C offset in H+ vs H-) of the heatwave in H+. Overall, H+ had a stronger immediate negative effect on cladoceran biomass (LM, p<0.001, Appendix Table A3), while recovery was not affected by any of the treatments. Results of the linear models testing for resistance (i.e., biomass difference between t_1_ and t_2_) and resilience (i.e., biomass difference between t_3_ and t_4_) are presented in Table S4.

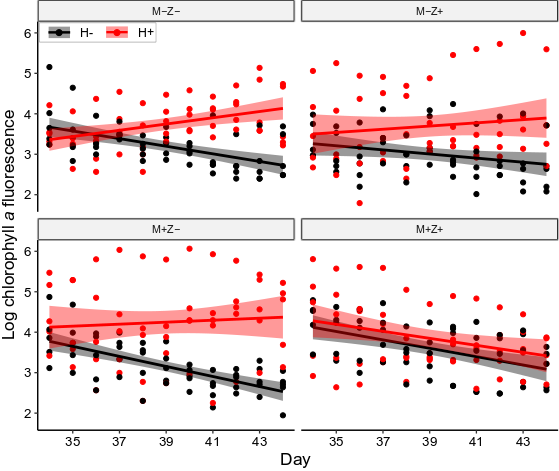

### **Figure S6** – Pattern of daily in vivo fluorescence of chlorophyll *a* (proxy for phytoplankton biomass) in the second part of the recovery period, grouped according to the dispersal treatments (microorganism: M- and M+; zooplankton: Z- and Z+) and coloured according to the heatwave treatment (H- and H+). N = 5 for all treatment combinations per day. Solid lines represent fitted linear models (error bands: 95% confidence intervals) to visualise temporal trends. Based on linear mixed-effects models, H+ drove an increase of phytoplankton over time relative to the H- treatment (LMEM: significant day × H+, p < 0.05, Table 2).

### **Table S4** – Summary statistics (estimated parameters, standard error and p values) of multiple linear regression models testing the ratio of zooplankton carbon mass to POC (<100-µm fraction) at the four focal sampling points. Significant results (p<0.05) are highlighted with bold letters while marginally significant ones (p<0.1) with italics. Note that for t_2_, t_3_ and t_4_, the analyses were performed on log-transformed data (to normalise model residuals). Treatment labels: H+: heatwave, M+: dispersal of microorganisms, Z+: dispersal of zooplankton.

|  | **t_1_** | | |  | **t_2_ (log)** | | |  | **t_3_ (log)** | | |  | **t_4_ (log)** | | |
| --- | --- | --- | --- | --- | --- | --- | --- | --- | --- | --- | --- | --- | --- | --- | --- |
|  | est. | SE | *p* |  | est. | SE | *p* |  | est. | SE | *p* |  | est. | SE | *p* |
| intercept | **0.53** | **0.10** | **<0.001** |  | 0.11 | 0.16 | 0.493 |  | **-1.04** | **0.23** | **<0.001** |  | **-0.63** | **0.18** | **0.001** |
| H+ | 0.02 | 0.14 | 0.895 |  | **-0.73** | **0.23** | **0.004** |  | **-0.68** | **0.32** | **0.035** |  | **-0.54** | **0.26** | **0.041** |
| M+Z- | -0.14 | 0.14 | 0.318 |  | -0.19 | 0.23 | 0.415 |  | 0.24 | 0.32 | 0.438 |  | -0.11 | 0.26 | 0.680 |
| M-Z+ | -0.13 | 0.14 | 0.334 |  | -0.32 | 0.23 | 0.181 |  | -0.07 | 0.32 | 0.832 |  | 0.36 | 0.26 | 0.170 |
| M+Z+ | 0.15 | 0.14 | 0.281 |  | -0.39 | 0.23 | 0.104 |  | 0.04 | 0.32 | 0.902 |  | -0.29 | 0.26 | 0.275 |
| H+ × M+Z- | 0.24 | 0.19 | 0.224 |  | -0.18 | 0.33 | 0.580 |  | *-0.79* | *0.45* | *0.087* |  | -0.31 | 0.36 | 0.394 |
| H+ × M-Z+ | 0.11 | 0.19 | 0.582 |  | 0.08 | 0.33 | 0.809 |  | -0.31 | 0.45 | 0.487 |  | **-0.84** | **0.36** | **0.027** |
| H+ × M+Z+ | -0.20 | 0.19 | 0.314 |  | 0.26 | 0.33 | 0.428 |  | 0.05 | 0.45 | 0.918 |  | 0.14 | 0.36 | 0.711 |
| **R^2^** | 0.03 | | |  | 0.47 | | |  | 0.47 | | |  | 0.53 | | |

### **Table S5** – Dominant phytoplankton taxa at the three focal sampling dates based on average biomass (calculated for N = 24 mesocosms per each date).

| **Rank** | **t_1_** | **t_2_** | **t_4_** |
| --- | --- | --- | --- |
| **1.** | *Chromulina* (Ø 5-10 µm) | *Nitzschia* (25-50 µm) | *Nitzschia* (25-50 µm) |
| **2.** | *Navicula* (> 50 µm) | *Euglena* | *Scenedesmus* gr. *Acutodesmus* |
| **3.** | *Chromulina* (Ø 10-15 µm) | *Ankyra* | Dinophyceae (Ø 10-15 µm) |
| **4.** | Dinophyceae (Ø 20-25 µm) | *Navicula* (> 50 µm) | Ulothrichales sp.1 |
| **5.** | Dinophyceae (Ø 10-15 µm) | *Cymbella* (< 50 µm) | *Mougeotia* |
| **6.** | *Lobomonas* | *Phacus* | unicellular coccal green (Ø 2.5 µm) |
| **7.** | *Chlamydomonas* | *Nitzschia* (> 50 µm) | *Monoraphidium minutum* |
| **8.** | green flagellates (Ø 6x5 µm) | *Chromulina* (Ø 5-10 µm) | *Tabellaria flocculosa* |
| **9.** | *Chrysochromulina* | *Lepocinclis* | *Navicula* (> 50 µm) |
| **10.** | *Gymnodinium* sp.1 | *Nephrocytium* | *Chromulina* (Ø 5-10 µm) |
| **11.** | *Gymnodinium* sp.2 | *Fragilaria* (> 50 µm) | *Oedogonium* |
| **12.** | *Nitzschia* (> 50 µm) | green flagellates (Ø 6x5 µm) | *Ankyra* |
| **13.** | *Ankyra* | *Navicula* (< 50 µm) | Ulothrichales sp.2 |
| **14.** | *Nitzschia* (25-50 µm) | *Mougeotia* | Volvocales |
| **15.** | *Cymbella* (< 50 µm) | Volvocales | *Gomphonema* (> 50 µm) |
| **16.** | Cymbella (> 50 µm) | colonial Chlorococcales | *Cryptomonas* (< 20 µm) |
| **17.** | Navicula (< 50 µm) | *Mallomonas* | *Navicula* (< 50 µm) |
| **18.** | *Mougeotia* | *Hyalotheca dissiliens v. minor* | *Gomphonema* (< 50 µm) |
| **19.** | Volvocales | *Cryptomonas* (20-40 µm) | *Gymnodinium* sp.1 |
| **20.** | *Caloneis* | Centrales (> 20 µm) | *Hyalotheca dissiliens v. minor* |

### **Table S6** – Results of SIMPER analysis listing taxa with the highest contribution to the overall dissimilarity among those treatments, which had a significant effect on community composition after the heatwave (t_2_) and in the recovery phase (t_4_) based on PERMANOVA (Table 2). For phytoplankton and zooplankton, the five and three most influential taxa (i.e., based on their contributions) are listed along with their mean biomasses (expressed in carbon mass, µg L^-1^) in treatments. P values indicate significant associations with treatments, originating from permutation tests (n = 1000).

| **Comparison** | **Overall dissimilarity** | **Taxa** | **Contribution %** | **Cumulative %** | **Mean biomass in ‘-‘** | **Mean biomass in ‘+’** | ***p*** |
| --- | --- | --- | --- | --- | --- | --- | --- |
| **t_2_  - Zooplankton** | | | | | | | |
| H- vs H+ | 31.7 | ***Daphnia* cf. *longispina*** | 21.8 | 68.8 | 26.2 | 13.3 | **<0.001** |
|  |  | *Cyclops abyssorum* | 3.4 | 79.6 | 7.2 | 6.9 | 1.000 |
|  |  | ***Bosmina longispina*** | 2.4 | 87.0 | 2.0 | 0.6 | **0.001** |
| **t_4_ – Phytoplankton** | | | | | | | |
| H- vs H+ | 65.8 | *Nitzschia* sp. | 21.3 | 32.4 | 14.5 | 13.7 | 0.537 |
|  |  | *Scenedesmus* gr. *Acutodesmus* | 10.4 | 48.2 | 0.3 | 11.2 | **0.019** |
|  |  | Ulotrichales | 7.1 | 59.0 | 0.8 | 3.2 | **0.031** |
|  |  | *Mougeotia* | 4.0 | 65.1 | 2.5 | 1.5 | 0.663 |
|  |  | unidentified green (2.5µm) | 4.0 | 71.1 | 1.3 | 2.6 | 0.478 |
| M- vs M+ | 65.4 | *Nitzschia* sp. | 22.4 | 34.2 | 13.1 | 15.2 | 0.138 |
|  |  | *Scenedesmus* gr. *Acutodesmus* | 9.8 | 49.3 | 0.9 | 10.6 | 0.401 |
|  |  | Ulotrichales | 6.6 | 59.4 | 3.6 | 0.4 | 0.288 |
|  |  | *Mougeotia* | 4.8 | 66.8 | 3.0 | 1.1 | **0.006** |
|  |  | unidentified green (2.5µm) | 3.8 | 72.7 | 1.7 | 2.3 | 0.905 |
| **t_4_ – Zooplankton** | | | | | | | |
| H- vs H+ | 38.1 | ***Daphnia* cf. *longispina*** | 23.1 | 60.8 | 24.3 | 11.7 | **0.001** |
|  |  | copepod nauplii | 4.0 | 71.2 | 2.3 | 3.4 | 0.057 |
|  |  | *Cyclops abyssorum* | 3.3 | 79.9 | 5.4 | 5.8 | 0.998 |

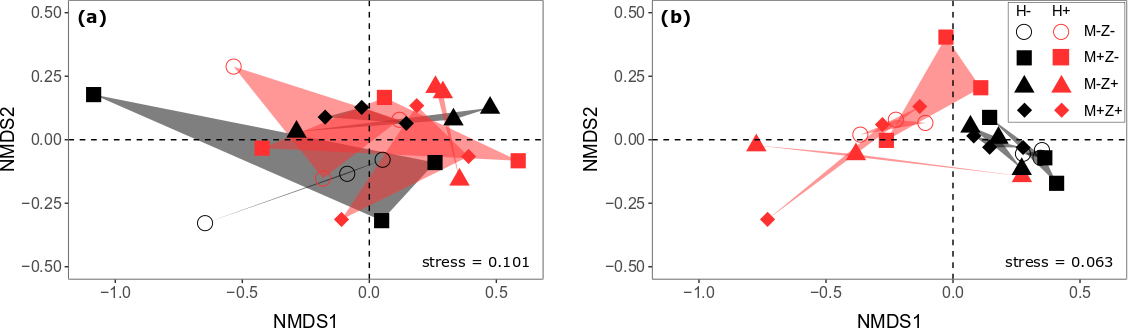

### **Figure S7** – NMDS plots illustrating the effects of heatwave (H-, H+) and dispersal treatments (microorganisms: M-, M+, zooplankton: Z-, Z+) on (a) phytoplankton and (b) zooplankton community composition (based on biomasses) at the end of the heatwave (t_2_). Zooplankton community composition was significantly influenced by H+ (PERMANOVA: p<0.001), while we observed no treatment effect on phytoplankton. Results of PERMANOVAs are presented in Table 2.
